# Extensive genome-wide association analyses identify genotype-by-environment interactions of growth traits in Simmental cattle

**DOI:** 10.1101/2020.01.09.900902

**Authors:** Camila U. Braz, Troy N. Rowan, Robert D. Schnabel, Jared E. Decker

## Abstract

**Background:** Understanding the genetic basis of genotype-by-environment interactions (GxE) is crucial to understand environmental adaptation in mammals and improve the sustainability of agricultural production. In addition, GxE information could also be useful to predict the vulnerability of populations to climate change.

**Results:** Here, we present an extensive study investigating the interaction of genome-wide SNP markers with a vast assortment of environmental variables and searching for SNPs controlling phenotypic variance (vQTL) using a large beef cattle dataset. We showed that GxE contribute 10%, 4%, and 3% of the phenotypic variance of birth weight, weaning weight, and yearling weight, respectively. GxE genome-wide association analysis (GWAA) detected a large number of GxE loci affecting growth traits, which the traditional GWAA did not detect, showing that functional loci may have non-additive genetic effects between genotype classes regardless of differences in genotypic means. We also showed that variance-heterogeneity GWAA can detect loci enriched with GxE effects without requiring prior knowledge of the interacting environmental factors. Functional annotation and pathway analysis of GxE genes revealed biological mechanisms by which cattle respond to changes in their environment, such as neural signaling, metabolic, hypoxia-induced, and immune system pathways. Knowledge of these pathways will be important as climate change becomes a burden on animal health and productivity. In addition, ecoregion-specific GxE SNPs detected in this study may play a crucial role in identifying resilient and adapted beef cattle across divergent environments.

**Conclusions:** We detected novel trait associations with large GxE effects for birth weight, weaning weight, and yearling weight. Functional annotation and pathway analysis uncovered genomic regions involved in response to environmental stimuli. We unraveled the relevance and complexity of the genetic basis of GxE underlying growth traits, providing new insights into how different environmental conditions interact with specific genes influencing adaptation and productivity in beef cattle and potentially across mammals

## Background

Genotype-by-environment interactions (GxE) refer to a variable response of genotypes across environments, resulting in different trait values [1,2]. GxE is a central issue in genetics, evolution, and ecology [3], and cattle represent a unique opportunity to address these questions due to a large number of genotyped animals [4] reared over a wide range of climatic and topographic regions. GxE may contribute to poor adaptation, which negatively affects the profitability and sustainability of the beef cattle production systems [1,5]. Studies using the relationship between breeding values and the environment have reported the existence of GxE for body weight in beef cattle [6,7]; however, limited research has focused on understanding the genetic loci of GxE underlying such traits. This knowledge could be used to better drive selection and mating decisions [8] as well as in statistical models incorporating biological information to accomplish more accurate genomic predictions [9]. In addition, GxE information could also be useful to predict the vulnerability of populations to climate change [3] and to understand environmental adaptation in other mammalian species.

Here, we present a comprehensive study of GxE for birth weight (BW), weaning weight (WW), and yearling weight (YW) using SNP marker information and over 13,000 Simmental cattle. First, we estimated the contribution of GxE variance on the total phenotypic variances using a restricted maximum likelihood (REML) approach. Then, we searched for SNPs enriched for GxE using four linear mixed model approaches: i) direct GxE genome-wide association analyses (GxE GWAA) using continuous environmental variables; ii) direct GxE GWAA using discrete environmental variables (United States ecoregions); iii) variance-heterogeneity genome-wide association analyses (vGWAA), which enable identification of interactions, using phenotypes adjusted for additive relationship matrix or additive, dominance and epistatic relationship matrices; and iv) meta-analysis of ecoregion-specific GWAA. Lastly, we performed enrichment analyses using GxE candidate genes to investigate the biological mechanisms by which environmental factors modulate such phenotypes. To our knowledge, this is the most comprehensive study investigating the interactions of genome-wide SNP markers with a vast assortment of environmental variables and searching for SNPs controlling phenotypic variance using a large beef cattle dataset. Our results revealed novel trait associations and alternative biological mechanisms involved in shaping the total phenotypic variance of growth traits providing new insights into how the environment influences growth traits and adaptation in beef cattle and potentially across mammals.

## Results

### Environmental variance component estimates

Phenotypes were pre-adjusted for fixed effect of sex and random effect of contemporary groups. The amount of BW, WW, and YW variation explained by contemporary group effects was 19%, 53%, and 61%, respectively, showing that the effect of farm management on the animal body weight increases with age. After adjusting the phenotypes, we estimated the contribution of GxE variance on the total phenotypic variances using additive relationship matrix calculated based on SNP marker information, and United States ecoregions as environmental factor. Ecoregions were determined based on the combination of 30-year normal values of mean temperature, precipitation, and elevation [10]. Each 1 km square of the United States was then assigned to one of 9 resulting ecoregions. We found that GxE variances contribute in 10.1%, 3.8%, and 2.8% to the BW, WW, and YW total phenotypic variances, respectively, suggesting that there is a decrease in the effect of GxE on the animal body weight throughout life. These results indicate an inverse relationship between farm management and environmental stress and that management practices (e.g. season of birth, forage quality and quantity, feed supplementation, health programs) are more relevant for later weights than climate environmental factors. One possible explanation for these findings is that birth weight reflects intrauterine growth which is highly affected by environmental conditions [11]; however, as the animal ages physiological, morphological, endocrine, cellular and molecular mechanisms are altered in order to cope up with environmental challenges [12]. These results also suggest that as management inputs are decreased, animals become more sensitive to environmental stress, which has ramifications for animal performance as we seek to decrease inputs to make beef production more sustainable.

### Genotype-by-environment GWAA

The GxE GWAA for BW, WW, and YW were performed by fitting univariate and multivariate linear mixed models considering continuous or discrete environmental variables. The continuous environmental variables used were minimum, mean, maximum, and mean dew point temperatures; elevation; precipitation; and minimum and maximum vapor pressure deficit, separately (Additional file 1: Figures S1–S8). Discrete environmental variables included United States ecoregions (Desert & Arid Prairie, Southeast, High Plains, Forested Mountains, Fescue Belt, and Upper Midwest & Northeast – Fig. 1F), which are combinations of mean temperature, precipitation, and elevation 30-year normal values (Additional file 1: Figures S9–S14). Descriptive information for growth traits and environmental variables are included in Additional file 2: Table S1. Univariate and multivariate models using both continuous and ecoregions environmental variables detected a total of 2,319 GxE SNPs (*P* < 1e-5, Table 1, Additional files 3 and 4). Only 25% of the GxE SNPs detected by the ecoregion GxE GWAA were also identified by the continuous GxE GWAA. These results indicate that the use of ecoregion information as environmental variable allowed us to capture GxE SNPs interacting with region-specific stressors that were not directly measured (e.g. forage differences, water availability, pathogens) and the combination of multiple stressors.

**Table 1.** Number of significant SNPs (*P* < 1e-05) and magnitude of the allele substitution effects by environmental variables/factors for each trait.

| Env | Birth Weight |  | Weaning Weight |  | Yearling Weight |  | Mv | Total |
| --- | --- | --- | --- | --- | --- | --- | --- | --- |
|  | N | ASE | N | ASE | N | ASE | N | N |
| Elev | 142 | -0.001–0.001 | 28 | -0.008–0.002 | 15 | -0.005–0.006 | 50 | 216 |
| Precip | 338 | -0.001–0.002 | 4 | -0.004– -0.004 | 1 | -0.02 | 12 | 349 |
| MT | 555 | -0.170–0.201 | 1 | 0.49 | 4 | -1.240–1.704 | 69 | 611 |
| MinT | 756 | -0.159–0.212 | 6 | -0.868–0.439 | - | - | 88 | 823 |
| MaxT | 408 | -0.175–0.207 | 1 | 0.43 | 4 | -1.267–1.848 | 23 | 427 |
| MDPT | 941 | -0.178–0.168 | 4 | -0.516–0.409 | 1 | -1.11 | 94 | 1018 |
| MinVPD | 152 | -1.399–1.831 | 2 | -3.254– -3.242 | 1 | -6.62 | 10 | 165 |
| MaxVPD | 224 | -0.213–0.262 | 10 | -0.546–0.556 | 1 | 1.08 | 17 | 236 |
| SE | 54 | -2.078–1.649 | - | - | 2 | -11.120– -10.500 | 7 | 59 |
| HP | 40 | -1.381–1.132 | 16 | -5.353–3.930 | 4 | -22.900–6.722 | 23 | 63 |
| FM | 90 | -3.135–1.215 | 31 | -6.785–4.737 | 49 | -9.397–17.690 | 148 | 221 |
| FB | 47 | -2.145–0.953 | 1 | -2.94 | 1 | 18.53 | 11 | 56 |
| UN | 24 | -0.709–0.872 | - | - | 1 | 8.18 | 10 | 33 |
| DA | 125 | -8.243–6.043 | 8 | -11.570–6.456 | 6 | -16.450–23.620 | * | 125 |
| Total | 1923 |  | 101 |  | 78 |  | 353 | 2319 |

Results of the genotype-by-environment genome-wide Association Analysis using environmental variables (Env), such as elevation (Elev), precipitation (Precip), mean temperature (MT), minimum temperature (MinT), maximum temperature (MaxT), mean dew point temperature (MDPT), minimum vapor pressure deficit (MinVPD), and maximum vapor pressure deficit (MaxVPD), and using the United States ecoregions (Southeast [SE], High Plains [HP], Forested Mountains [FM], Fescue Belt [FB], Upper Midwest & Northeast [UN], and Desert & Arid Prairie [DA]) as environmental factors (Env. Factor) in univariate (Birth Weight, Weaning Weight, and Yearling Weight) and multivariate (Mv) linear mixed models. The number of significant SNPs (N) at the significant threshold of $P < 1e-05$ and the magnitude of the allele substitution effects (ASE, kg) were also provided.

**Fig. 1.**
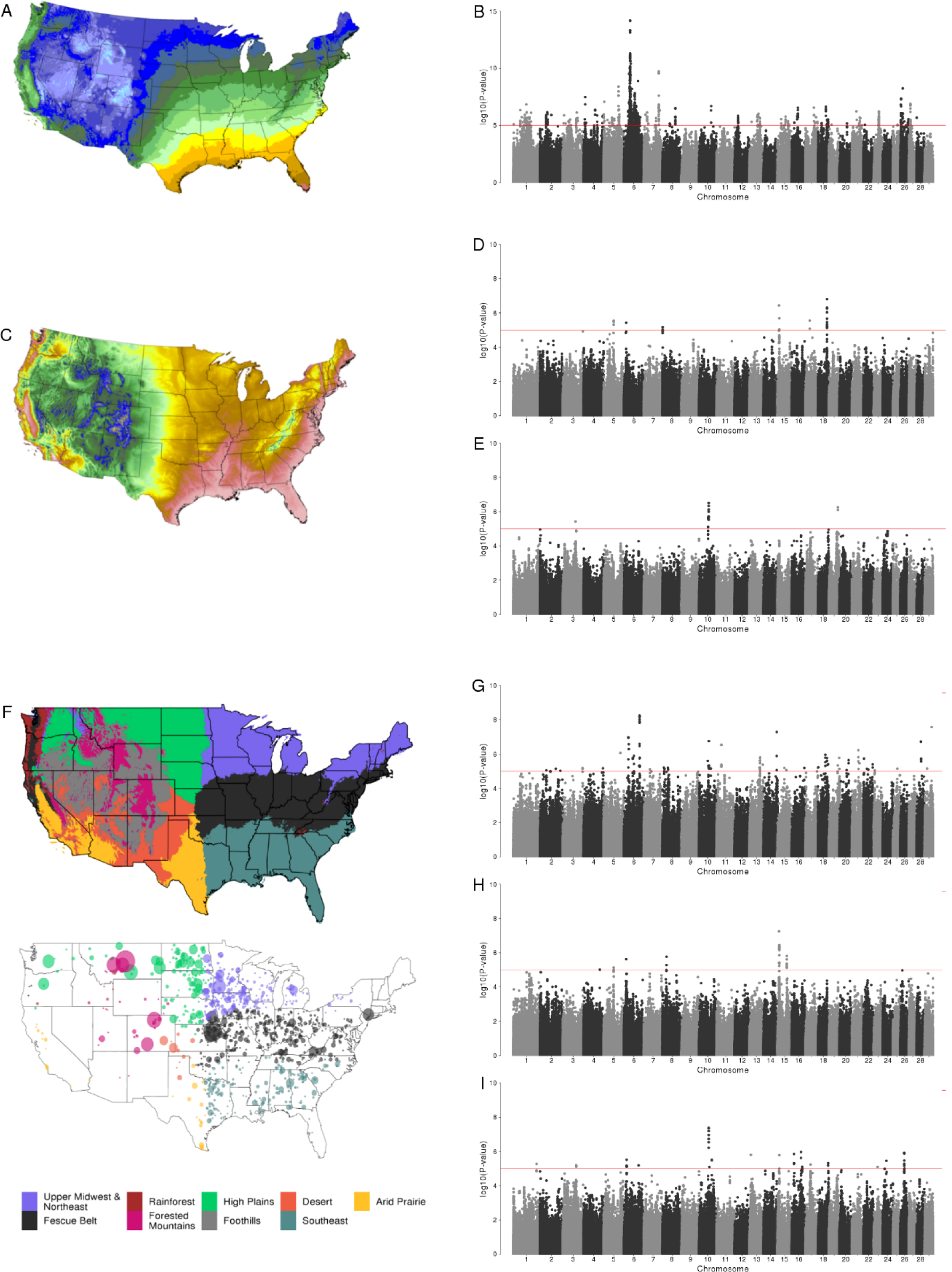
Genotype-by-environment interaction genome-wide association analyses (GxE GWAA) for growth traits in Simmental cattle. (*A*) Mean dew point temperature average annual over the most recent three full decades covering the conterminous United States. (*B*) Manhattan plot of GxE GWAA of mean dew point temperature for birth weight. (*C*) Elevation of the conterminous United States. (*D*) Manhattan plot of GxE GWAA of elevation for weaning weight. (*E*) Manhattan plot of GxE GWAA of elevation for yearling weight. (*F*) Boundaries for ecoregion assignments in the United States; (*Top*) United States partitioned into nine ecoregions based on similar topographic and environmental conditions; (*Bottom*) location of beef farms for which data was retrieved. (G) Manhattan plot of GxE GWAA of Desert & Arid Prairie ecoregion for birth weight. (*H*) Manhattan plot of GxE GWAA of Forested Mountains ecoregion for weaning weight. (*I*) Manhattan plot of GxE GWAA of Forested Mountains ecoregion for yearling weight. In Manhattan plots, horizontal red line indicates a significant threshold (*P* < 1e-5). Environmental continuous variables were drawn from the PRISM climate dataset (http://prism.oregonstate.edu). The United States was partitioned into nine regions using k-means clustering.

There was no overlap between the GxE SNPs identified for all traits using univariate models, however, multivariate models detected several GxE SNPs associated with BW, WW, or YW (*P* < 1e-5, Table 1). Multivariate analyses were able to detect GxE SNPs that were not identifiable by standard univariate analyses, indicating advantages of incorporating a multivariate statistical approach to GxE genome-wide association studies of correlated traits. Consistent with our GxE variance component results, a larger number of GxE SNPs was detected for BW than for WW or YW. BW was highly affected by all continuous environmental variables and ecoregions analyzed (Table 1). However, mean dew point temperature was the environmental variable that most interacted with SNPs for BW (*P* < 1e-5, Fig. 1B), and consequently, the ecoregion that identified more GxE SNPs for BW was the Desert & Arid Prairie (*P* < 1e-5, Fig. 1G). The Desert & Arid Prairie ecoregion has both high temperatures and many locations with low absolute humidity measured by dew point temperature (Fig. 1A, Additional file 2: Table S1), which in concert both affect BW. The effect of higher temperature on calf birth weight may be due to the reduced feed intake or the decreased uterine blood flow and placental function often found in heat-stressed cows, factors that contribute to impaired cow-to-fetal nutrient exchange, affecting fetal growth [13–15]. For WW and YW, elevation (Fig. 1C) was found to be the continuous environmental variable that most interacted with SNPs (*P* < 1e-5, Fig. 1 D and E respectively). In agreement, Forested Mountains, the ecoregion with the highest elevation (Additional file 2: Table S1), showed more SNP-by-environment interaction for WW and YW (*P* < 1e-5, Fig. 1 *H* and *I* respectively), compared with other ecoregions. Higher altitude regions are associated with reduced atmospheric oxygen (hypoxia), which causes pulmonary hypertension and vasoconstriction leading to right-side heart failure in cattle [16]. This condition is known as brisket disease and accounts for significant losses in growth and reproductive performance, being one of the top causes of mortality in cattle residing at high altitudes [17].

Only one GxE SNP (*rs133638007*, chromosome 18, 61,721,281 bp) was detected interacting with two different ecoregions. The *rs133638007* T allele increase BW in 1.06 kg in the Desert & Arid Prairie ecoregion, on the other hand, results from the multivariate analysis showed that the *rs133638007* T allele decrease BW by 0.62 kg in Forested Mountains ecoregion. Interestingly, this SNP is located close to the gene *VSTM1*, a member of the leukocyte receptor complex, which is involved in immune responses [18]. In addition, the fact that only one GxE SNP was identified for two different ecoregions indicates that the loci interacting with abiotic and biotic stressors are different across divergent environments, and consequently, the ecoregions have specific genetic architecture involved in GxE.

We explored 10 kb and 100 kb sequence windows that flanked all significant GxE SNPs (*P* < 1e-5) to scan for genes in their vicinity and to identify possible regulatory elements (Additional file 5). Using these GxE gene sets, we performed functional enrichment analysis based on Gene Ontology (GO) and KEGG pathways to get insights into biological mechanisms by which beef cattle respond to changes in their environment. Response to stimulus was the most statistically significant GO term, and its direct descendants GO terms, cellular response to stimulus and response to stress, were also enriched (Additional file 2: Table S2). Additionally, many biological processes involved in metabolism, development, localization, gene expression, transport, signaling, and cell adhesion were enriched. Enriched pathways were related to metabolism; cell proliferation and differentiation; immune and inflammatory responses; hypoxia-induced processes; and mechanisms of neurotransmission in central nervous system (Additional file 2: Table S3). In addition, circadian entrainment, a pathway involved in environmental adaptation to natural periodic changes, including light/dark, temperature, humidity, social activity, and several other cycles [19], was enriched. All GO terms and pathways enriched and their respective GxE genes are available in Additional files 6 and 7, respectively.

As the genetic architecture of GxE appeared to be region-specific, we investigated whether they affect the same or different biological functions across ecoregions. To do so, we calculated the percentage of candidate GxE genes for each ecoregion that participate in each biological processes (GO) or pathways enriched (Fig. 2 and 3, Additional file 1: Figures S15 and S16). Even though the ecoregions exhibit mostly unique GxE QTL, the candidate genes associated with these QTLs point toward the same biological processes and pathways, however in different magnitudes. The GxE genes involved in response to stimulus were identified in all ecoregions. In the Southeast, 55.8% of the GxE genes detected play roles in the response to stimulus process, suggesting that the combined effects of high temperature and humidity pose a major challenge by affecting how the animal adjusts to external changes and maintains homeostasis. This will become a greater burden on animal health and productivity as the climate continues to warm. Indeed, studies have shown that heat and humidity stress disrupt homeostasis and result in compromised health, growth, fertility, and milk production in several species, including cattle, pig, sheep, mice and humans [12,20–23].

**Fig. 2.**
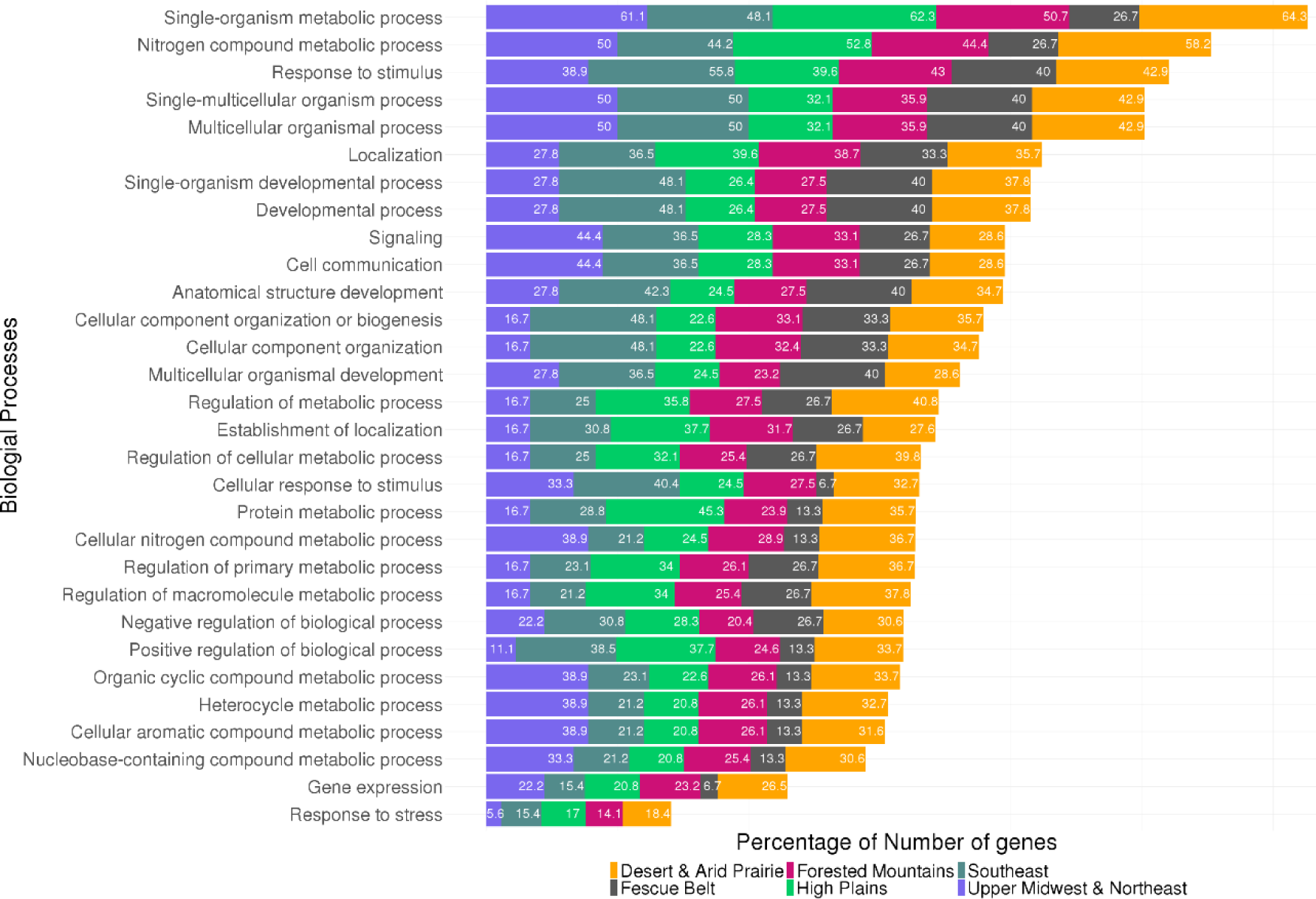
Percentage of candidate genes with genotype-by-environment effects for each ecoregion that contribute to the enriched gene ontology (GO) terms. Genes were located 100 kb from significant SNPs (*P* < 1e-5).

**Fig. 3.**
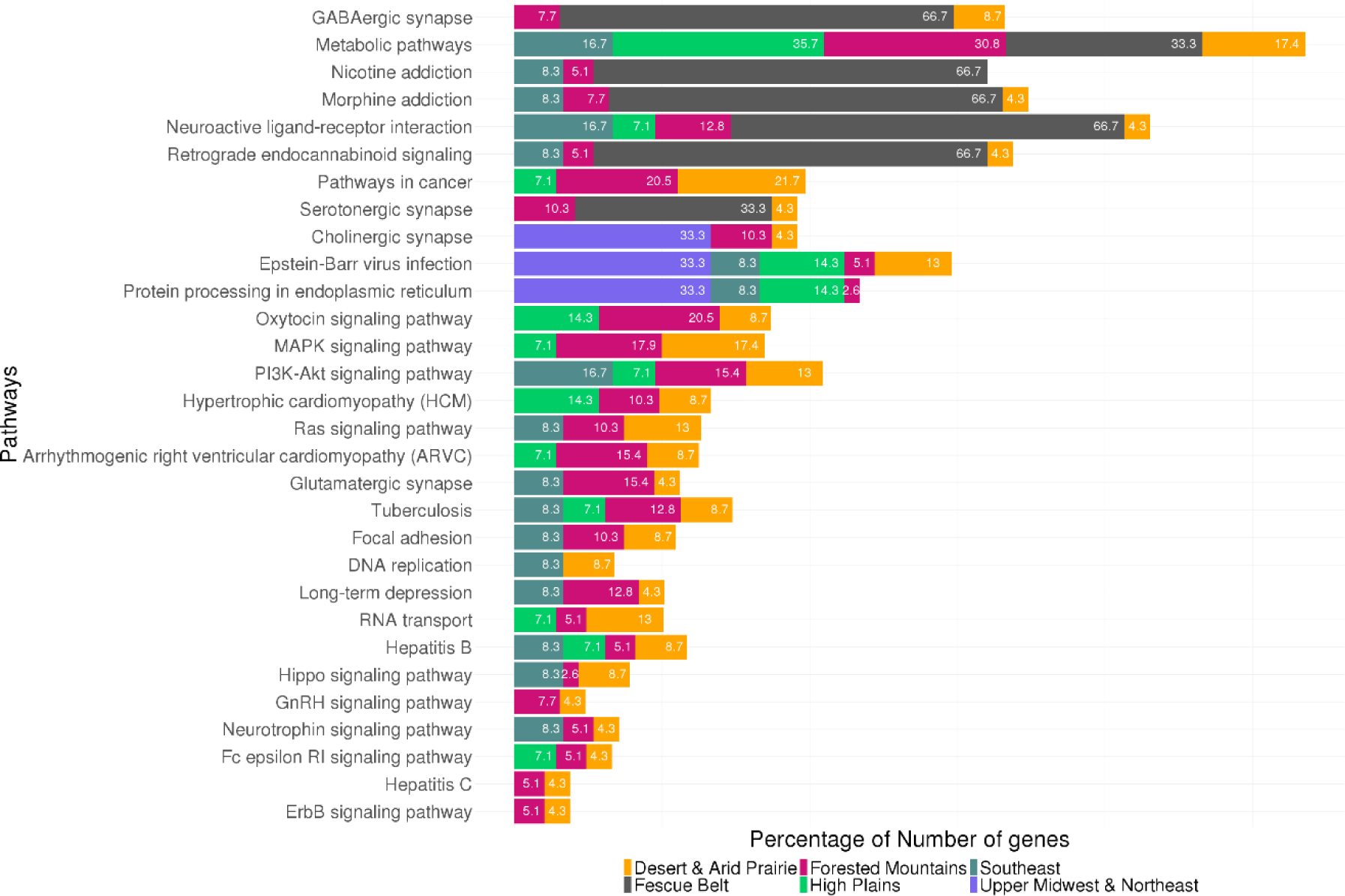
Percentage of candidate genes with genotype-by-environment effects for each ecoregion that contribute to the enriched biological pathways. Genes were located 100 kb from significant SNPs (*P* < 1e-5).

Cardiomyopathy related pathways were enriched predominantly in GxE genes interacting with ecoregions in the Western United States, as High Plains, Forested Mountains and Desert & Arid Prairie, ecoregions with higher elevation. Such pathways were identified likely due to the hypoxia condition exposure, which cause pulmonary arterial hyperplasia resulting in right ventricular hypertrophy and congestive/dilatory cardiac failure in cattle living at high altitude [17]. Pathways involved in neural development and neurotransmission processes were also enriched in GxE genes. Although these pathways were identified as being affected by GxE genes interacting with all ecoregions, they seem to be more affected in animals raised in the Fescue Belt and Southeast ecoregions. This may be due to the consumption of tall fescue, a grass commonly used in grazing cattle systems in the Southeastern United States [24], which usually contains an endophytic fungus that produces ergot alkaloids [25]. Ergot alkaloids have vasoconstrictive effects and interact with GABAergic, glutamatergic and serotonergic receptors affecting multiple aspects of animal’s physiology regulated by these neurotransmitters [26–28]. Animals consuming the endophyte-infected tall fescue forage often develop fescue toxicosis, which is characterized by reduced feed intake, weight gain, and blood flow, as well as elevated body temperature associated with heat stress [28–31]. Interestingly, the majority of GxE genes for the Fescue Belt and Southeast ecoregions were also found interacting with temperature. Numerous GxE genes involved in neural signaling were also detected in the Forested Mountains ecoregion. Hypoxic environments may affect cells of the nervous system as well. [32] reported that neuroactive ligand-receptor interaction participates in the regulation of neural stem cells behavior in mice under hypoxia. Pathways involved in hormone-releasing (oxytocin and GnRH signaling) related to acute stress in cattle [33] were also identified for high elevation ecoregions.

GxE genes of all ecoregions drove enrichments in biological pathways related to immune and inflammatory responses (Cytokine-cytokine receptor interaction, Fc epsilon RI signaling, Epstein-Barr virus infection, Tuberculosis, and Hepatitis), except Fescue Belt. This result is unexpected since studies have observed that fescue toxicosis causes hypoprolactemia leading to decreases in immune responses [34,35]. However, our results indicate that this effect may be due to the environmental condition of Fescue Belt ecoregion affecting the activity of GABA and serotonin. Such neurotransmitters are involved in the regulation of prolactin secretion [36,37] and, consequently, may affect the immune system. Many of these immune GxE genes have been associated with diseases caused by pathogens leading to reduced productivity in cattle [38–48]. This result is in agreement that heterogeneity in the immune response to infectious diseases across populations living in different environmental conditions is under genetic control [49]. In addition, pathogen resistance increases host fitness and has driven the evolution of plant and animals through time [50].

GxE genes also participate in pathways involved in the regulation of cell functions including proliferation, growth, gene expression, differentiation, cell survival, and apoptosis, which are crucial processes to proper animal growth and development [51]. Such pathways (PI3K-Akt signaling, Ras signaling, MAPK signaling, Hippo signaling, Focal adhesion, and ErbB signaling) were identified by GxE genes from all ecoregions. However, we identified more GxE genes involved in such pathways in the Southeast, Desert & Arid Prairie, Forested Mountains and High Plains ecoregions.

The majority of GxE SNP associations were identified for BW in this study. Studies have shown that loci identified in GWAA of BW could either represent direct effects of the fetal genotype, indirect effects of maternal genotype (acting via the intrauterine environment), or some combination of the two [52,53]. According to [54] the uterine environment may even have a greater influence on fetal growth and birth weight than the parental genome. Thus, we investigated whether GxE SNPs affecting BW could be reflecting maternal uterine environmental conditions. To test this, we adjusted BW by including random maternal effects in the model (MAM) and compared the results with those from the GxE GWAA without maternal effects (MA). Results showed that 876 SNPs (out of 1923) were not associated with GxE effects when accounting for the effect of dam, suggesting that those SNPs are interacting with the uterine environment. Candidate maternally-influenced GxE genes (Additional file 8) participate, in particular, in metabolic, insulin signaling, starch and sucrose metabolism, PI3K-Akt signaling and cGMP-PKG signaling pathways; and response to stimulus, blastocyst development, and regulation of signaling processes. These findings support previous investigations suggesting that detrimental environmental factors experienced by the cow during gestation may influence cow-to-fetal nutrient exchange limiting fetal growth and development [13–15,54–56]. Several maternally-influenced GxE genes identified in this study have been previously reported as maternal genes influencing embryonic development and/or response to environmental stress in cows and also in other mammals [57–66]. These results demonstrate that the bovine is an important model to understand the interaction between the physical environment, uterus, and fetus, and could be used in future research.

### Variance-heterogeneity GWAA

The vGWAA identify differences in variance, rather than differences in mean, between genotype classes, called vQTLs, which can be explained by GxE or GxG (epistatic) interactions [67]. Therefore, we performed a REML analysis to estimate how much of the BW, WW, and YW variance is explained by additive, dominance and epistasis effects. We fitted models (M) using only additive relationship matrix (MA) or using additive, dominance, and epistasis relationship matrices (MADE), see Additional file 2: Table S4. Overall, the heritabilities were greater than the ratio of the variance explained by the non-additive genetic effects to the phenotypic variances. Proportions of dominance variances in WW and YW were low (2% and 5%, respectively), whereas no dominance variance was estimated for BW. Epistatic relationship matrix accounted for 25%, 12% and 11% of the phenotypic variances for BW, WW, and YW, respectively, suggesting that epistasis has a substantial contribution to phenotypic variation for body weight traits in Simmental cattle. Thus, to search for vQTLs we used squared normalized residuals (z^2^), which is a measure of phenotypic variation [68], as dependent variables fitting univariate and multivariate models. Using z^2^ from the MA, we expected to detect loci with both GxE and GxG effects, since this phenotype was adjusted only for the additive relationship matrix. However, using z^2^ from the MADE, we should detect exclusively GxE loci as the phenotype was adjusted for additive, dominance and epistatic relationship matrices. In total, 44 SNPs with effect on z^2^ (*P* < 1e-5) were identified on chromosomes 3, 5, 8, 10, 11, 13, 14, 16, 17, 18, 21, 27, and 29 (Additional file 1: Figures S17 and S18, Additional file 2: Table S5). These 44 SNPs tag 24 putative vQTLs (grouped by LD patterns), with seven, four, and eleven vQTLs for univariate BW, WW, and YW, respectively. The multivariate analysis identified five vQTLs, of which two vQTLs (chromosome 8 at 4 Mb; and chromosome 29 at 39 Mb) were not identified by the univariate analyses.

Nine vQTLs were only significant using z^2^ from MA (three vQTLs for univariate BW, three vQTLs for univariate WW, two vQTLs for univariate YW, and one vQTL using multivariate analysis). Among these vQTLs likely involved in GxG, one vQTL for BW is located on chromosome 14 at 23 Mb and resides near or within the genes *LYN, RPS20, MOS, PLAG1*, and *CHCHD7*. This region has been reported to have epistatic effects on several chromosomes affecting post-weaning weight in beef cattle [69]. These effects could be due to the gene *PLAG1* which encodes a transcription factor that regulates the expression of several genes involved in a variety of cellular processes [70], and has been previously associated with stature and growth traits in many beef cattle breeds [71–73]. In addition, we detected a GxG vQTL on chromosome 3 at 106 Mb within *TRIT1*, a gene that was also reported to harbor a vQTL involved in epistatic interactions in humans [74].

Using z^2^ from MADE, 15 vQTLs were detected (four vQTLs for univariate BW; one vQTL for univariate WW; nine vQTLs for univariate YW; and four vQTLs using multivariate analyses, of which three vQTLs were also identified by the univariate analyses), which likely identify GxE. Based on GO information, genes near or within these vQTLs are involved in response to stimulus (*NOS1, FBLN1, YWHAH*, and *DEPDC5*), inflammation and immunity mechanisms (*LYZL1, KSR2, FBXO21, IL18RAP, XCL2, XCL1, TTYH1, LENG8, LENG9, CDC42EP5, LAIR1, OSCAR*, and *TARM1*), neurogenesis (*ZHX2*, and *NOS1*), regulation of transcription (*ZHX2, YWHAH* and *CNOT3*), developmental process (*FBXW8*, and *CNOT3*), transmembrane transport (SLC9A2, *SLC9A4, SLC5A1, SLC5A4*, and *TTYH1*), neurotransmission regulation (*SHANK2*), and cell proliferation (*DPT, RPS9*, and *TACC1*). Some of those genes were previously reported to be involved in environmental adaptation in cattle [*XCL2* [75], *SLC9A4* [76], *ZMAT4* [77]], and heat tolerance in buffalo [*XCL1* [78]] and dairy cattle [*SHANK2* [79]].

Using z^2^ as dependent variables in the analyses allowed the estimation of the proportion of residual variance that is explained by the SNPs in GEMMA (PVE). For BW z^2^, WW z^2^ and YW z^2^, SNPs explained 4.9% (SD = 0.8), 4.7% (SD = 0.8), and 5.9% (SD = 1.1), respectively, showing that residual variance is heritable and could be changed by selection, supporting previous studies [80–82]. Residual variability may reflect differences between animals in uniformity for a certain trait, so the reduction of variance could improve economic merit [80]. Rönnegard et al. [81] reported advantages in using breeding values explaining differences in residual variance (vEBV) for milk yield uniformity in Holstein cattle. According to [80] the reduction of phenotypic variance by selecting for reduced residual variance could be more beneficial for low heritability traits.

### Meta-analysis of ecoregion-specific GWAA

We performed a meta-analysis of ecoregion-specific GWAA (GWAA for each ecoregion separately) to look for significant differences (Cochran’s Q statistics) in effect size of SNPs between the ecoregions, which are interpreted as GxE [83]. In addition, we calculated the m-value statistic for the significant SNPs (Cochran’s Q statistic’s *P*-value < 1e-5), which is the posterior probability of an effect being present in an ecoregion given the observations from all other ecoregions [83] allowing us to identify which ecoregions a given variant has an effect. Southeast and Desert & Arid Prairie ecoregions were not included in the meta-analysis due to their small sample sizes. Meta-analysis of ecoregion-specific GWAA detected 12 GxE SNPs for BW located on chromosome 6 (Additional file 1: Figure S19A); two GxE SNPs for WW on chromosome 15 (Additional file 1: Figure S19B); and two GxE SNPs for YW, one located on chromosome 13 and one on chromosome 15 (Additional file 1: Figure S19C, Additional file 9). Analyzing the PM-plot (*P*-values against m-values) and PB-plot (*P*-values against beta-values) of the GxE SNPs detected by the meta-analysis, we observed that some GxE SNPs on chromosome 6 at 30 Mb have an effect on animal’s BW from Forested Mountain and Upper Midwest & Northeast ecoregions, however with different certainty (Additional file 1: Figures S20A and S21A). For WW and YW, despite that the analysis had shown a significant heterogeneity between the ecoregions, they did not show different posterior probabilities of an effect in any ecoregion (m-value), (Additional file 1: Figure S20B and C, respectively). However, large differences in the SNP effect size (beta) were noticed between ecoregions (Additional file 1: Figure S21B and C, respectively).

### Traditional genome-wide association analyses

We also performed traditional GWAA using univariate and multivariate linear mixed models in order to compare with the GxE GWAA results. We detected major QTLs on chromosomes 6 (26-48 Mb), 7 (88-90 Mb), 14 (15-30 Mb), and 20 (41-78 Mb) for univariate BW, WW and YW (Additional file 1: Figure S22A-C and Additional file 10). In addition, significant SNPs (*P* < 1e-5) were identified on chromosomes 3 (53.6 Mb) and 17 (73 Mb) for univariate BW; and on chromosomes 1 (81.1-81.3 Mb), 2 (116.1 Mb), and 5 (96.7 Mb) for univariate WW. The GWAA using multivariate model detected QTL also identified in the univariate GWAA on chromosomes 5, 6, 7, 14, 17, and 20 in addition to on chromosomes 4 (24.5 Mb), 9 (57.9 Mb), 13 (63.3 Mb), 19 (50.5 Mb), and 26 (45.4 Mb) (Additional file 1: Figure S22D and Additional file 11). All genes detected by GWAA are included in the Additional file 12.

Out of 948 total significant SNPs using multivariate GWAA analysis for the growth traits, 531 SNPs were located on chromosome 6 (from 26.8 to 48.8 Mb). Likewise, several SNPs interacting with Forested Mountains ecoregion, elevation, precipitation and, mainly, with the temperatures were also detected in the same positions on chromosome 6. Given the importance of chromosome 6 underlying the growth traits, we repeated the multivariate GWAA including significant SNPs (e.g. *rs109849093, rs464458177, rs110305942, rs43459303, rs109278547*, and *rs137209027* located at 37,211,057; 37,234,136; 37,418,164; 37,695,352; 38,139,495; and 41,084,765 bp, respectively) as fixed effects in the model. In this way, if those SNPs are causal mutations accounting for QTL effect the loci association signals would be missed. Results showed that even including six significant SNPs (not in LD) simultaneously as fixed effects in the model, a small number of SNPs were still associated (six out of 531 SNPs), indicating that there are possibly multiple causal mutations in this region.

### Comparison of the GWA approaches

Comparing traditional GWAA with all GxE GWAA results, we verified that 312 SNPs with additive effects (QTL) also have GxE effects (mainly on chromosome 6, but also on chromosomes 7 and 20). Some SNPs showed GxE effects even greater than their additive effects on the traits. For example, the SNP *rs136523580* (chromosome 6 at 30,358,905) has an additive effect of −0.43 kg for BW in the total dataset, but its GxE effects is even more impactful (−0.67 kg) on the BW of animals from the Forested Mountains ecoregion. The SNP *rs109868461* (chromosome 6, at 36,902,704 bp) has additive effects of −0.69 kg for BW in the total dataset, however it has a GxE effect of 1.11 kg on the BW of animals born in the Southeast ecoregion. These findings show the importance of GxE effects on quantitative traits and the need for precision, ecoregion-specific genomic predictions to optimize animal performance.

Comparing vGWAA with the traditional GWAA results, only one vQTL, on chromosome 14 at 23.1-23.3 Mb harboring the genes *LYN, MOS, RPS20, PLAG1*, and *CHCHD7*, was also found as a direct effect QTL for BW, meaning that this locus affects both mean and variance of BW in this population. Five vQTLs were also identified by the GxE GWAA, which were found to interact with multiple continuous environmental variables as well as interacting with ecoregions. These results agree with [84] in suggesting that vQTL effects are much more enriched with GxE effects than additive effects. However, GxE GWAA detected a greater number of GxE loci when compared with the vGWAA results.

Only few GxE SNPs were detected using the meta-analysis of ecoregion-specific GWAA (Cochran’s Q statistic’s *P*-value < 1e-5), which were also identified by the GxE GWAA. In addition, using the meta-analysis approach the Southeast and Desert & Arid Prairie ecoregions could not be included in the analysis due to their small sample sizes, suggesting that this was not an optimal approach to detect GxE associations for our dataset. However, creating PM-plots (contrast between environment-specific *P*-value and posterior probability of an effect, m-value)[85], and PB-plots (environment-specific *P*-value and SNP effect size) helped interpret results of our GxE GWAA and vGWAA. For significant GxE SNPs detected by the GxE GWAA, there were clear differences in the SNP effect sizes between the significant ecoregion and the remaining ecoregions (Additional file 1: Figures S23 and S24). For SNPs with significant variance-heterogeneity effects (detected by vGWAA), patterns were not as consistent. While some vQTLs had differences in mean effects, others did not (Additional file 1: Figures S25 and S26).

## Discussion

This GxE study provides a more comprehensive understanding of how GxE modulate growth traits in beef cattle and possibly in other mammalian species. We observed that GxE explain substantial proportions of growth traits phenotypic variances, which is likely due to a great number of GxE SNPs with large effects affecting such traits. Detection of large GxE SNP effects indicates that there is considerable opportunity to improve environmental resistance and performance in cattle. Our study also revealed the first evidence of vQTLs controlling phenotypic variation in body weights due to GxG or GxE effects in beef cattle. In addition, we uncovered several biological pathways affected by GxE that likely drive environmental adaptation.

One of our major findings is that the ecoregions showed specific GxE with large effects on the traits. In addition, ecoregion GxE GWAA detected GxE loci that were not identified by the continuous GxE GWAA. One possible explanation for these results is that when analyzing ecoregions, we take into account not only climate and topography, but also unmeasured variables like forage quality and local pathogens, as well as the combination of them. This information allowed us to identify environmentally sensitive genotypes. It indicates that the same trait can be influenced by different genes in the various ecoregions which may affect particular physiological and behavioral functions leading to local adaptation. This assumption is supported by the GO and pathways enrichment analysis of the GxE genes identified for each ecoregion showing that the diverse ecoregion environment affects different biological mechanisms and in different magnitude. We also found that major additive QTLs also carry GxE effects on growth traits and that some of their GxE effects are greater or in opposite directions when compared with their additive effects. Additionally, novel GxE associations were detected in multiple genomic regions. These findings support the importance of creating ecoregion-specific genomic predictions to identify local resilient or adapted animals, using the GxE SNPs detected in this study to assist selection decisions which would lead to an increase in animal performance within an ecoregion.

Some vQTLs were also identified by the GxE GWAA, indicating that vGWAA can detect loci enriched with GxE effects without requiring prior knowledge of the interacting environmental factors. Such GxE loci showed interaction with several continuous and discrete (ecoregions) environmental variables in the GxE GWAA, suggesting that vGWAA may capture GxE loci that interact with multiple environmental variables, possibly even those not included in the GxE GWAA model (e.g. water availability, pasture condition, pathogens). According to [86] when multiple interacting factors induce variance heterogeneity, the power of identification of any single one of them or all together may be lower than the power of the variance heterogeneity test, which may explain why vGWAA identified GxE loci that were not detected by direct GxE GWAA. We also observed that some vQTLs have differences in means across ecoregions (Additional file 1: Figure S27), however, not all do. This suggests that some vQTLs could be the difference between robust genotypes (little variation, flat reaction norms) across environments while other genotypes at the same locus are more variable across environments, indicating that selection for low variance vQTL genotypes may provide sustainability advantages. Many genes harboring the vQTL identified here were previously reported to be involved in environmental adaptation in cattle [75–79] supporting that phenotypic variability may be an adaptive evolutionary solution to environmental changes [87]. However, our results support [88] which reported that when an environmental factor of interest has been measured on all the genotyped individuals, the direct GxE GWAA will be able to detect a larger number of GxE loci.

We found that environmental conditions modulate growth traits through genes that participate in a variety of biological mechanisms that are activated to help the body cope with external stimuli and return to or maintain homeostasis [89,90]. Among them, several pathways involved in neural signaling and development were detected, which play crucial roles in the regulation of response to stress [91,92], body temperature, vasoconstriction [93,94], and circadian rhythms [95]. Specifically, for animals living in higher altitude ecoregions, we identified pathways related to hypoxia response and secretion of hormones [33]. Other environmental pathways included immune functions reflecting the involvement of innate and adaptive immune systems in the activation of host defense reactions to deal with pathogens [94]. We also identified many metabolic processes that may be involved in acute stress responses and thermoregulation [12,33,94]. These biological mechanisms are affected differently by the diverse ecoregion environments and have been reported to be involved in resilience [12,92,94], and adaptation processes [10,16,96–98], revealing the role that GxE plays in adaptation and evolution at the level of the genome in cattle and potentially across mammals. It is important to understand such biological mechanisms since the inability of an animal to cope with its environment results in failure to produce optimally leading to lost potential profitability of livestock operations [12,17,33,99]. Further, these biological insights may help us understand how adaptation occurs in wild populations and the potential effect of climate change on animal physiological functions.

## Conclusions

In summary, GxE GWAA and vGWAA detected several loci affecting growth traits that the traditional GWAA did not, showing that functional loci may have non-additive genetic effects between genotype classes regardless of differences in genotypic means. We revealed biological mechanisms by which beef cattle respond to changes in their environment. Neural signaling, metabolic, hypoxia-induced, and immune system pathways were highlighted, indicating that climate change may become a burden on animal health and productivity. In addition, ecoregion-specific GxE SNPs detected in this study may play a crucial role in resilience and adaptation of beef cattle to divergent environments. Although conventional genomic selection has shown great promise in improving genetic gain, it only considers additive effects and, according to our findings, ecoregion-specific genomic predictions should be created to identify animals best suited for given environments. This could help producers in making optimal decisions given their geographical location. Our results revealed novel trait associations and alternative genetic mechanisms involved in shaping the total phenotypic variance of growth traits providing new insights into how the environment influences such traits and adaptation in beef cattle and other mammalian species.

## Methods

### Phenotype, genotype and environmental data

Data from the American Simmental Association males and females born between 1975 and 2016 and genotyped for genomic-enhanced EPDs were transferred to the University of Missouri. The traits studied were birth weight (BW), 205-d adjusted weaning weight (WW) and 365-d adjusted yearling weight (YW), with the traits also adjusted for age of dam as per breed association practice. The animals were assigned to contemporary groups (CG), defined by the combination of farm, season (spring vs. fall) and year of birth.

Phenotypes were pre-adjusted for fixed effect of sex estimated using ‘*--reml-est-fix*’ option, and random effect of contemporary groups (CG) was predicted by the BLUP method using ‘*--reml-pred-rand*’ flag while controlling for population stratification using single-trait animal model implemented in GCTA software [100]. CG were included in the univariate model as random effect since 19% of individuals analyzed belong to CG with only a single animal [101]. The random maternal permanent environmental effect was also included in the WW adjustment.

The DNA samples were genotyped with various low-density assays and were imputed to the combination of Illumina BovineHD (Illumina, San Diego, CA) and the GeneSeek Genomic Profiler F250 (GeneSeek, Lincoln, NE) according to [102]. Genotype filtering was performed to remove non-autosomal variants, as well as both variants and individuals with call rates less than 0.90, using PLINK 1.9 [103]. SNP positions were based on the ARS-UCD1.2 Bovine reference genome assembly [104]. The genotypes were then phased using Eagle 2.4 [105] and imputed using Minimac3 [106]. The final imputed genotype data contained a total of 835,947 bi-allelic variants with imputation accuracy of 99.6% [102]. Variants with minor allele frequency less than 0.01 were removed using PLINK 1.9 for further analysis, leaving 710,202, 709,054, and 706,771 SNP markers for 13,427, 11,847, and 8,546 animals for BW, WW and YW, respectively.

Eight continuous environmental variables were used for the GxE GWA analyses, including 30-year normal of minimum, mean, maximum and mean dew point temperatures; elevation; precipitation; and minimum and maximum vapor pressure deficit, which were drawn from the PRISM climate dataset [107]. The United States was partitioned into nine regions (Fig. 1F *Top*) based on similar topographic and environmental conditions (ecoregions), including mean temperature, elevation and precipitation information, using k-means clustering implemented in ‘*RStoolbox*’ R package [108,109]. The optimal ecoregions were identified using ‘*pamk*’ function from the R package ‘*fpc*’ [110]. The ecoregions were named Desert (D), High Plains (HP), Arid Prairie (AP), Forested Mountains (FM), Upper Midwest & Northeast (UN), Southeast (SE), Rainforest (R), Foothills (FH), and Fescue Belt (FB) [10]. Animals were assigned to the ecoregions based on the zip code of their breeder (Fig. 1F *Bottom*), and only ecoregions with more than 200 records were analyzed. Thus, animals located in the Rainforest ecoregion were removed from the dataset, and those that were reared in the Desert and Arid Prairie ecoregions were analyzed together (called DA) due to their similar environmental conditions. No animal record was assigned to the Foothills ecoregion. A complete description of the dataset, including weight records and environmental variable measurements is shown in Additional file 2: Table S2.

### Estimation of variance components

Variance components for pre-adjusted BW, WW, and YW were estimated using multi-component restricted maximum likelihood (REML) approach, implemented in GCTA software, in univariate linear mixed models that included only additive effects (MA), or additive, dominance and epistasis effects (MADE) as follows:

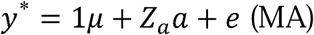

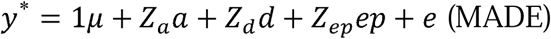

where *y*\* is the vector of pre-adjusted phenotypes; *µ* is the overall mean; *a* is the vector of random additive genetic effects; is *d* the vector of random dominance effects; *ep* is the vector of random epistatic effects; and is *e* the vector of random residuals. ***Z*** matrices are incidence matrices relating observations to animals. Assuming that 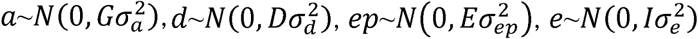, where 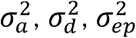, and 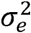 are the additive, dominance, epistatic, and residual variances, respectively. ***I*** is the identity matrix; and ***G, D***, and ***E*** are the additive, dominance and epistatic genetic relationship matrices, respectively. The genomic (***G***) and the dominance (***D***) relationship matrices were calculated using SNP marker information according to [111] as: 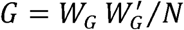, and 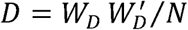, where 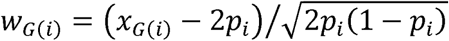 and, 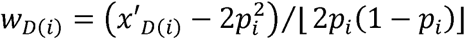 with *x*_*G*_ being 0, 1, or 2 and being 0, 2*p*, or (4*p* − 2) for genotypes AA, AB or BB, respectively; *p*_*i*_ is the allele frequency at SNP *i*; and *N* is the number of SNPs. According to Henderson [112] the epistatic relationship matrix (***E***) can be derived from the additive genomic relationship matrix as: *E*= *G*# *G*, where # denotes the Hadamard product operation.

The variance of GxE interaction was also estimated considering the ecoregions as an environmental factor and treated as random effect in the model in order to estimate the amount of phenotypes that are explained by the GxE interaction among the ecoregions, as implemented in GCTA.

### Genome-wide association analyses (GWAA)

Univariate linear mixed model analyses were performed for the pre-adjusted BW, WW or YW using GEMMA software [113] as follows:

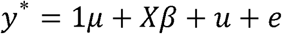

where *y*\* is an *n*-vector of pre-adjusted phenotypes; *µ* is the overall mean; *X* is the incidence matrix of genotypes; *β* is the genotype effects; *u* is an *n*-vector of random additive genetic effects; and *e* is an *n*-vector of random residual. Assuming that *u* ∼ *MVN*_*n*_(0, *GV*_*g*_) and *e* ∼ *MVN*_*n*_(0,*IV*_*e*_), ***G*** is the *n* × *n* genomic relationship matrix (GRM); *V*_g_ is the residual additive genetic variance; *V*_*e*_ is the residual variance component; ***I*** is an *n* × *n* identity matrix. *MVN*_*n*_ denotes the *n*-dimensional multivariate normal distribution. GRM were estimated using the standardized genotypes of 710,202, 709,054, and 706,771 SNP markers for 13,427, 11,847, and 8,546 animals for BW, WW and YW, respectively.

We also fitted a multivariate linear mixed model for the pre-adjusted BW, WW and YW using GEMMA [114] in the following form:

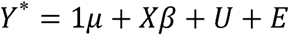

where ***Y***^*∗*^ is an *n* × *d* matrix of *d* pre-adjusted phenotypes for *n* individuals; *µ* is the overall mean; ***X*** is the incidence matrix of genotypes; *β* is a *d* vector of the genotype effects for the *d* phenotypes; ***U*** is an *n* × *d* matrix of random additive genetic effects; and ***E*** is an *n* × *d* matrix of random residuals. *U*∼ *MN*_*nxd*_ (*0, G, V*_*g*_) and *E*∼ *MN*_*nxd*_ (0, *I*_*nxn*_, *V*_*e*_), ***G*** is the *n* × *n* genomic relationship matrix (GRM); *V*_*g*_ is the *d* × *d* matrix of additive genetic variance component; *V*_*e*_ is a *d* × *d* matrix of the residual variance component; ***I*** is an *n* × *n* identity matrix. *MN*_*nxd*_ denotes the *n* × *d* matrix normal distribution with mean 0, row covariance matrix ***G*** or ***I*** (*n* × *n*), and column variance matrix *V*_*g*_ or *V*_*e*_ (*d* × *d*). For all GWA analyses performed, single-marker *P*-values were used to generate Manhattan plots using ‘*manhattan*’ function implemented in ‘*qqman*’ R package [115].

### Genotype-by-environment GWA analyses

The GxE GWAA were performed using GEMMA through two approaches: using eight continuous environmental variables (i.e. minimum, mean, maximum and mean dew point temperatures; elevation; precipitation; and minimum and maximum vapor pressure deficit) separately; and using the ecoregions as discrete environmental variable, where each ecoregion was compared against the total dataset using 0 and 1 dummy coding. In the GxE GWAA, for each SNP in turn, GEMMA fits a linear mixed model that controls both the SNP main effect and environmental main effect, while tested for the interaction effect and controlling for population stratification, evaluating the alternative hypothesis (H_1_: ≠ 0) against the null hypothesis (H_0_:*x*_j_*β*_*ij*_ = 0) for each interaction, therefore the resulting *P*-values correspond to the significance of the GxE interaction [71]. This model can be described as:

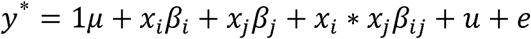

where y* is an *n*-vector of pre-adjusted phenotype; *x*_*i*_ is an *n*-vector of the genotypes of the SNP *i*; *β*_*i*_ is the effect size of the SNP *i*; *x*_*j*_ is an *n*-vector of environmental variable *j*; *β*_*j*_ is the fixed effect of the environmental variable; *β*_*ij*_ is the interaction effects between the genotypes of the SNP *i* and the environmental variable; *u*is an *n*-vector of random additive genetic effects; and *e* is an *n*-vector of random residual. GxE GWA analyses using multivariate linear mixed model were also performed.

### Variance-heterogeneity GWAA

We performed vGWAA to detect locus affecting the difference in variance between genotypes (vQTL). Residual effects (*e*) from the MA and MADE models were standardized to z score by the rank-based inverse-normal transformation and squared (z^2^), which is a measure of phenotypic variance [84,116]. The vGWAA were performed for BW z^2^, WW z^2^, and YW z^2^ using univariate and multivariate linear mixed models implemented in GEMMA software as denoted in “Genome-wide association analyses” section, where y* is an *n*-vector of z^2^; and *β*_*i*_ is the effect size of the SNP *i* on z^2^. Linkage disequilibrium (LD) analyses were performed using PLINK 1.9. All vQTL were defined by SNP markers with r^2^ greater than 0.6 [117].

### Meta-analysis of ecoregion-specific GWAA

In order to evaluate how significantly the effect size of SNPs (Cochran’s Q test) are between the ecoregions (indicative of GxE) [83] we combined the outputs from univariate within-ecoregion GWAA into a single meta-analysis for each phenotype using METASOFT software [118]. Only ecoregions with more than 1000 animals were analyzed. In addition, the statistic m-value for the significant SNPs (Cochran’s Q statistic’s *P*-value < 1e-5) were calculated, which is an estimative of the posterior probability of a locus having an effect in a particular ecoregion [119]. The results were visualized through PM-plots and PB-plots, in which *P*-values are simultaneously visualized with the m-values and beta-values at each tested locus, respectively.

### False discovery rate (FDR) estimation

FDR of the nominal significant threshold (*P* < 1e-5) was estimated for all analyses performed (Additional file 2: Table S6), based on [120], as:

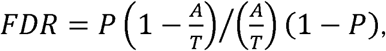

where *P* is the nominal significant threshold (e.g., 1e-5); *A* is the number of SNP that were significant at the nominal significant threshold; and *T* is the total number of SNPs tested. Although the true FDR cannot exceed 1, the estimated FDR can exceed 1 if the number of significant SNPs is smaller than expected by chance, meaning that all SNPs that met the nominal significant threshold are false positives [69].

### Functional enrichment analyses

We explored 10 kb and 100 kb sequence windows that flanked the significant SNPs (*P* < 1e-5) to scan for GxE genes located in their vicinity and to identify possible regulatory elements, respectively, based on the ARS-UCD1.2 bovine reference genomic positions. Enrichment analysis using the GxE genes based on Gene Ontology (GO) and KEGG pathways were performed using ShinyGO v0.60 [121], in which the analyses were adjusted for FDR of 5%. GO annotations were also retrieved from the AmiGO browser [122].

## Supplementary information

**Additional file 1: Supplementary Figures**. Supplementary Figures S1-S26. (DOCX 5.0 MB)

**Additional file 2: Supplementary Tables**. Supplementary Tables S1-S6. (DOCX 272.9 KB)

**Additional file 3:** Significant SNPs (*P* < 1e-5) using univariate GxE GWAA for Birth Weight, Weaning Weight and Yearling Weight. (XLSX 248.7 KB)

**Additional file 4:** Significant SNPs (*P* < 1e-5) using multivariate GxE GWAA for Birth Weight, Weaning Weight and Yearling Weight. (XLSX 40.7 KB)

**Additional file 5:** Candidate genes identified by GxE GWAA using 10 kb and 100 kb region flanking the significant SNPs. (XLSX 253.8 KB)

**Additional file 6:** Enrichment analyses based on Gene Ontology terms using GxE genes for 10 kb and 100 kb region flanking the significant SNPs. (XLSX 1.2 MB)

**Additional file 7:** Enrichment analyses based on KEGG pathways using GxE genes for 10 kb and 100 kb region flanking the significant SNPs. (XLSX 86.1 KB)

**Additional file 8:** Candidate maternally-influenced GxE genes using 10 kb and 100 kb region flanking the significant SNPs. (XLSX 77.3 KB)

**Additional file 9:** Significant SNPs (Cochran’s Q statistic’s *P*-value < 1e-5) using meta-analysis of ecoregion-specific GWAA for Birth Weight, Weaning Weight and Yearling Weight. (XLSX 5.4 KB)

**Additional file 10:** Significant SNPs (*P* < 1e-5) using univariate GWAA for Birth Weight, Weaning Weight and Yearling Weight. (XLSX 128.8 KB)

**Additional file 11:** Significant SNPs (*P* < 1e-5) using multivariate GWAA for Birth Weight, Weaning Weight and Yearling Weight. (XLSX 93.6 KB)

**Additional file 12:** Candidate genes identified by GWAA for Birth Weight, Weaning Weight and Yearling Weight. (XLSX 19.6 KB)

## Declarations

### Ethics approval and consent to participate

Animal Care and Use Committee approval was not required or obtained for data that were extracted from the existing American Simmental Association database.

### Consent for publication

Not applicable.

### Availability of data and materials

The raw data used in this research are available from the American Simmental Association under a Data Use Agreement, but are not publicly available. Derived data (analytical results) are however available as supplementary files associated with this publication.

### Author’s contributions

CUB and JED conceived the study. CUB analyzed the data and interpreted the results and wrote the manuscript. TNR and RDS managed data and imputed variants. JED assisted with analyses, preparation of the manuscript and interpretation of results. All authors read and approved the final manuscript.

## Acknowledgments

The authors would like to acknowledge the American Simmental Association for providing phenotypic and genotypic data. We also appreciate the farmers and ranchers who collected this data.

