## Additional File 1 for "Extensive genome-wide association analyses identify genotype-by-environment interactions of growth traits in Simmental cattle"

Supplementary Figures

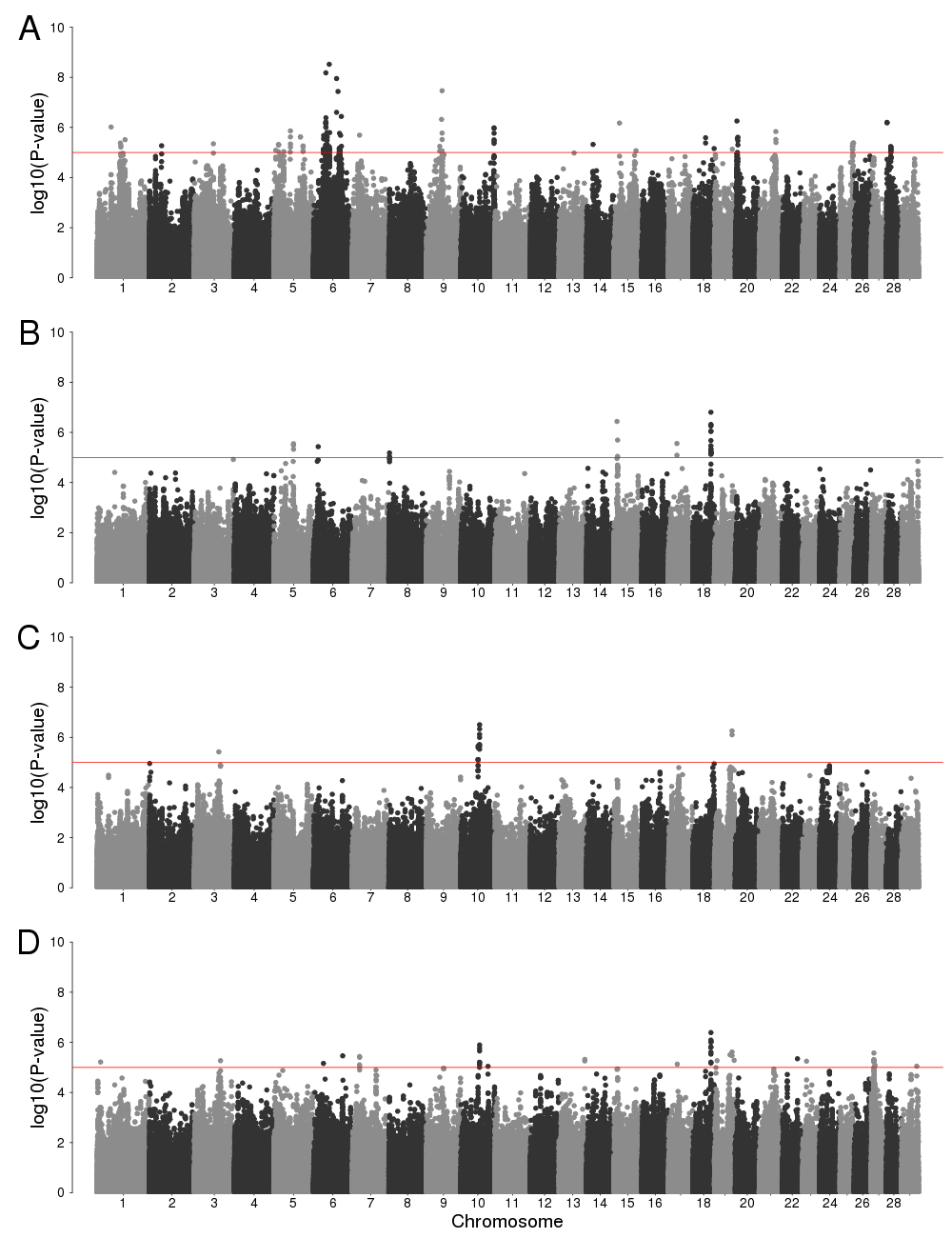
Figure S1 Manhattan plots of genotype-by-environment genome-wide association analysis using elevation as environmental variable for birth weight (*A*), weaning weight (*B*), yearling weight (*C*), and using multivariate analysis (*D*). Horizontal red line indicates a significant threshold (*P* < 1e-5).

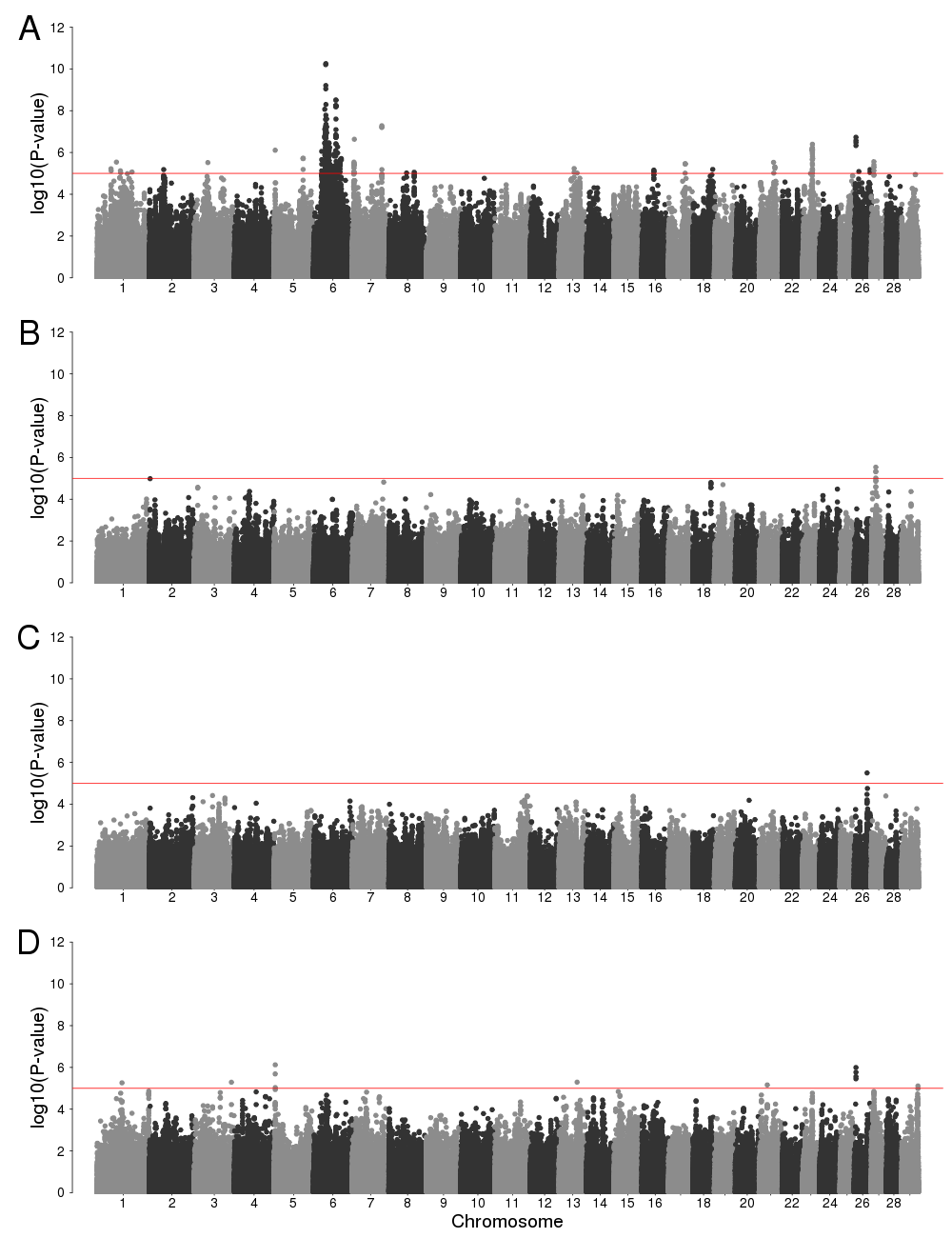
Figure S2 Manhattan plots of genotype-by-environment genome-wide association analysis using precipitation as environmental variable for birth weight (*A*), weaning weight (*B*), yearling weight (*C*), and using multivariate analysis (*D*). Horizontal red line indicates a significant threshold (*P* < 1e-5).

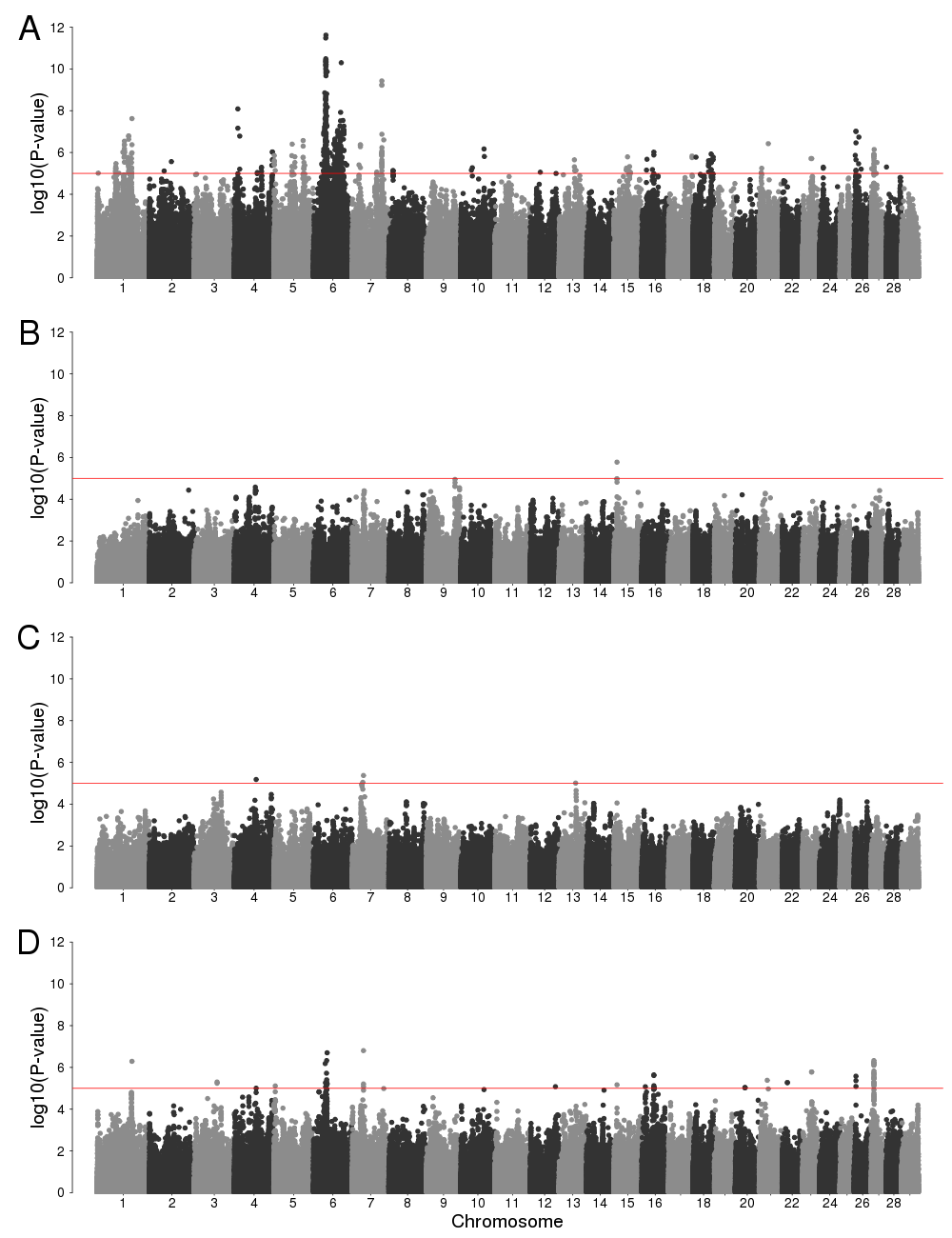
Figure S3 Manhattan plots of genotype-by-environment genome-wide association analysis using mean temperature as environmental variable for birth weight (*A*), weaning weight (*B*), yearling weight (*C*), and using multivariate analysis (*D*). Horizontal red line indicates a significant threshold (*P* < 1e-5).

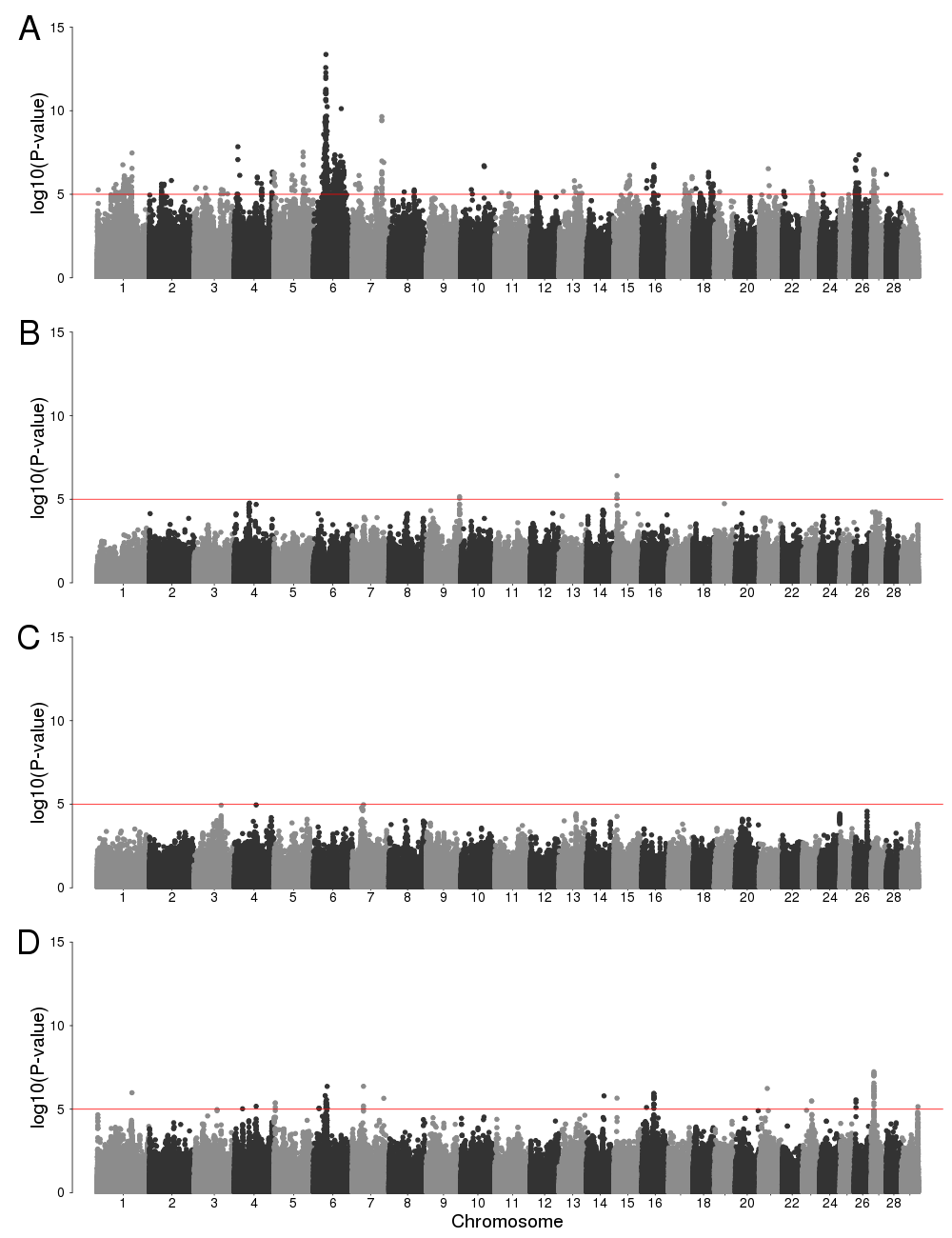
Figure S4 Manhattan plots of genotype-by-environment genome-wide association analysis using minimum temperature as environmental variable for birth weight (*A*), weaning weight (*B*), yearling weight (*C*), and using multivariate analysis (*D*). Horizontal red line indicates a significant threshold (*P* < 1e-5).

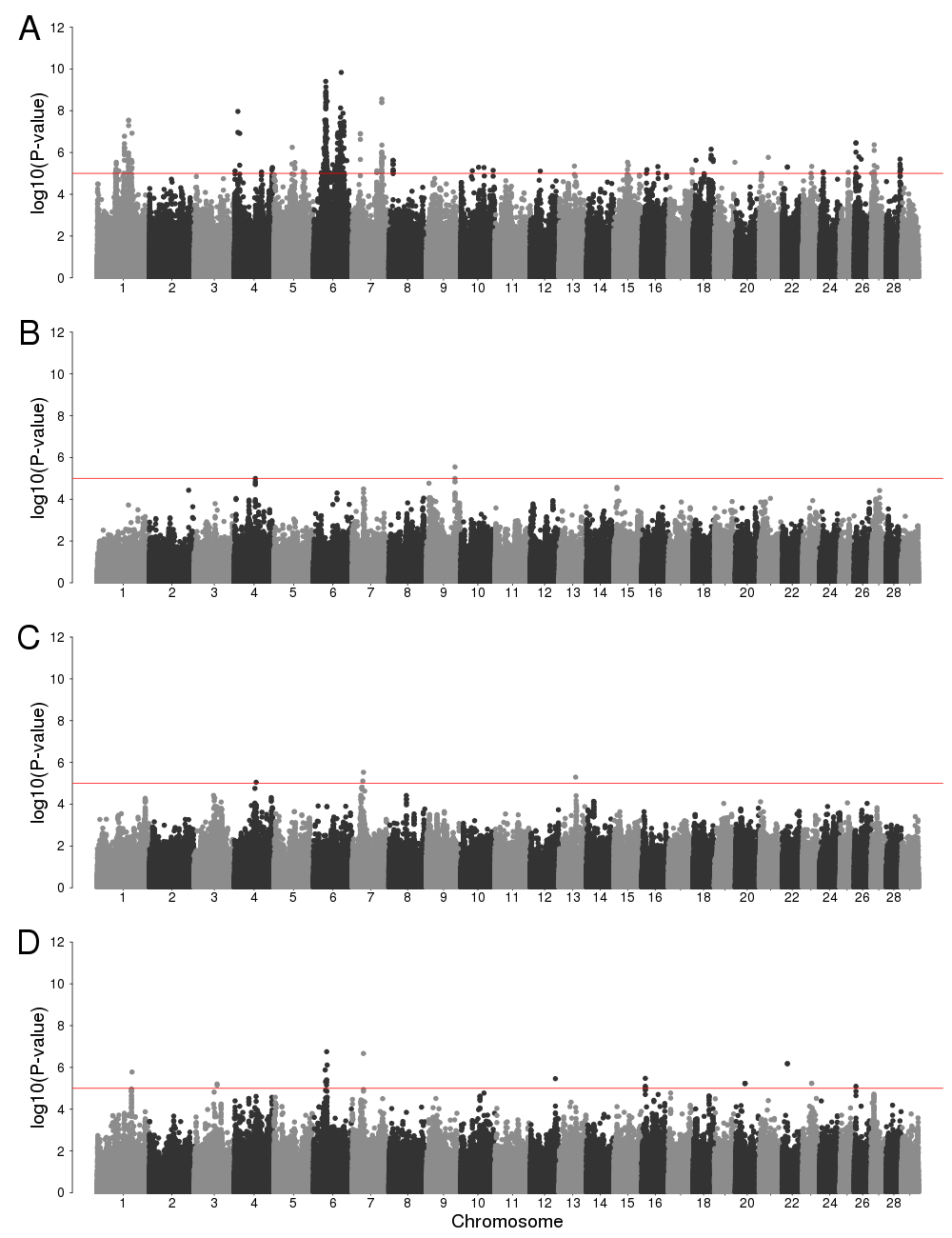
Figure S5 Manhattan plots of genotype-by-environment genome-wide association analysis using maximum temperature as environmental variable for birth weight (*A*), weaning weight (*B*), yearling weight (*C*), and using multivariate analysis (*D*). Horizontal red line indicates a significant threshold (*P* < 1e-5).

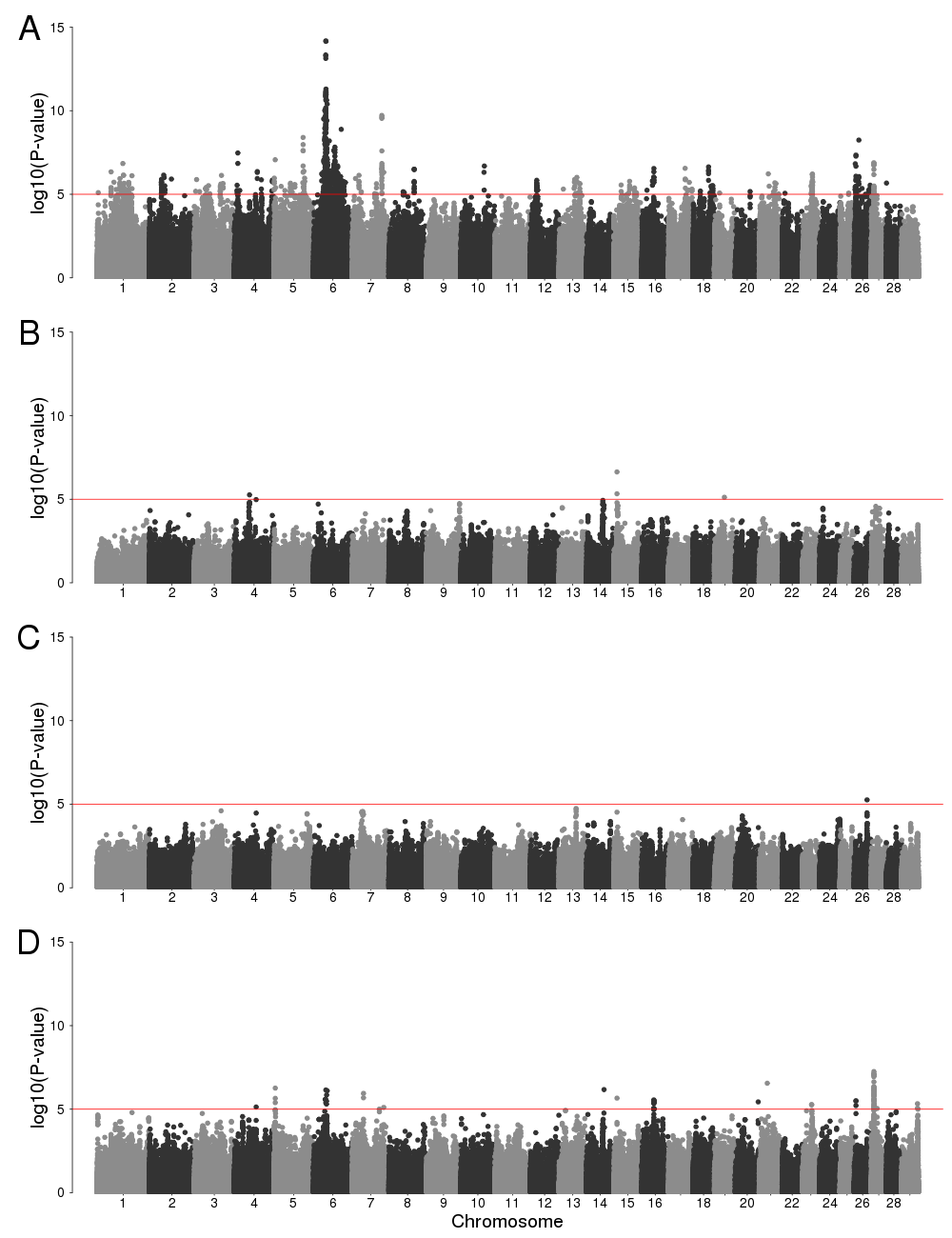
Figure S6 Manhattan plots of genotype-by-environment genome-wide association analysis using mean dew point temperature as environmental variable for birth weight (*A*), weaning weight (*B*), yearling weight (*C*), and using multivariate analysis (*D*). Horizontal red line indicates a significant threshold (*P* < 1e-5).

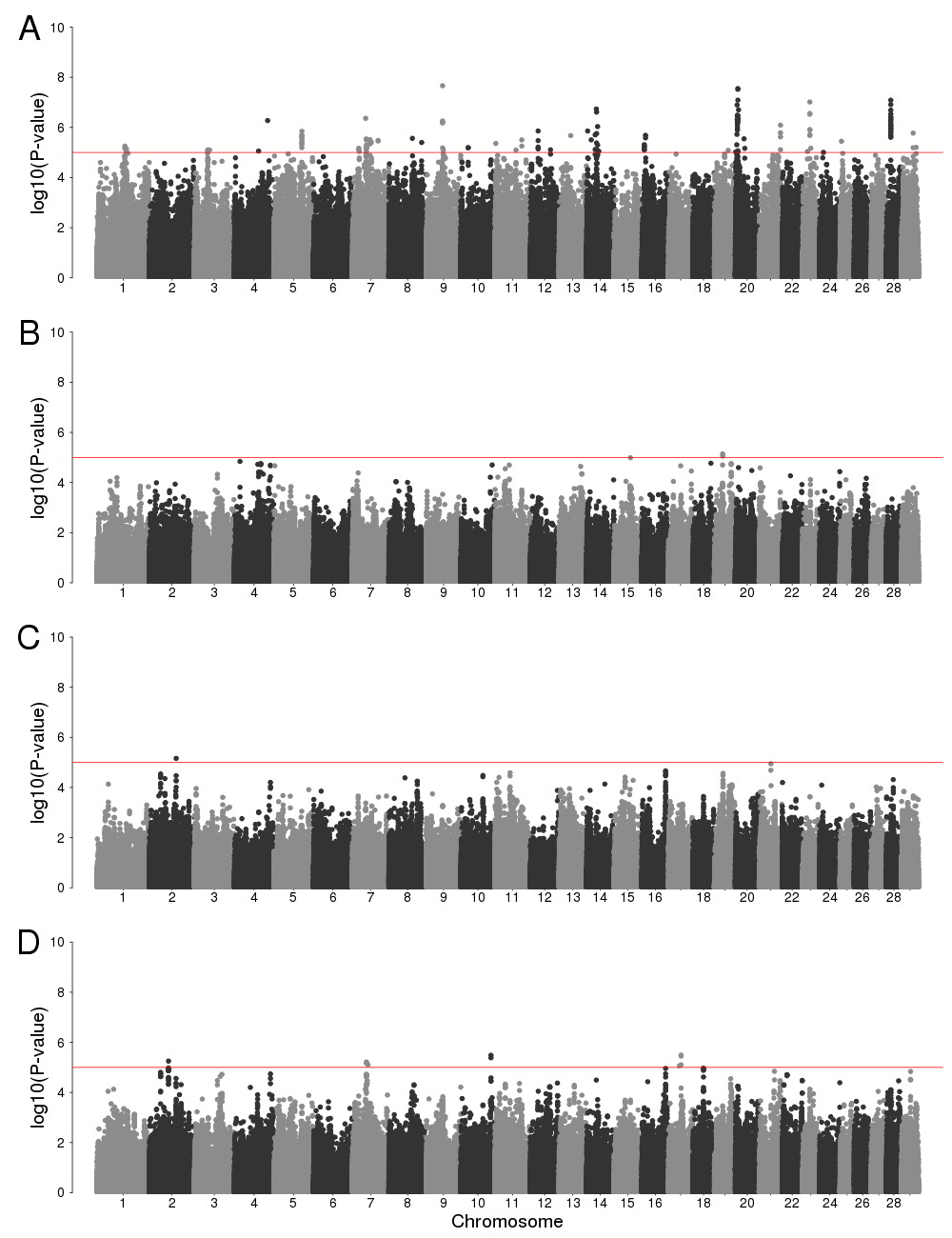
Figure S7 Manhattan plots of genotype-by-environment genome-wide association analysis using minimum vapor pressure deficit as environmental variable for birth weight (*A*), weaning weight (*B*), yearling weight (*C*), and using multivariate analysis (*D*). Horizontal red line indicates a significant threshold (*P* < 1e-5).

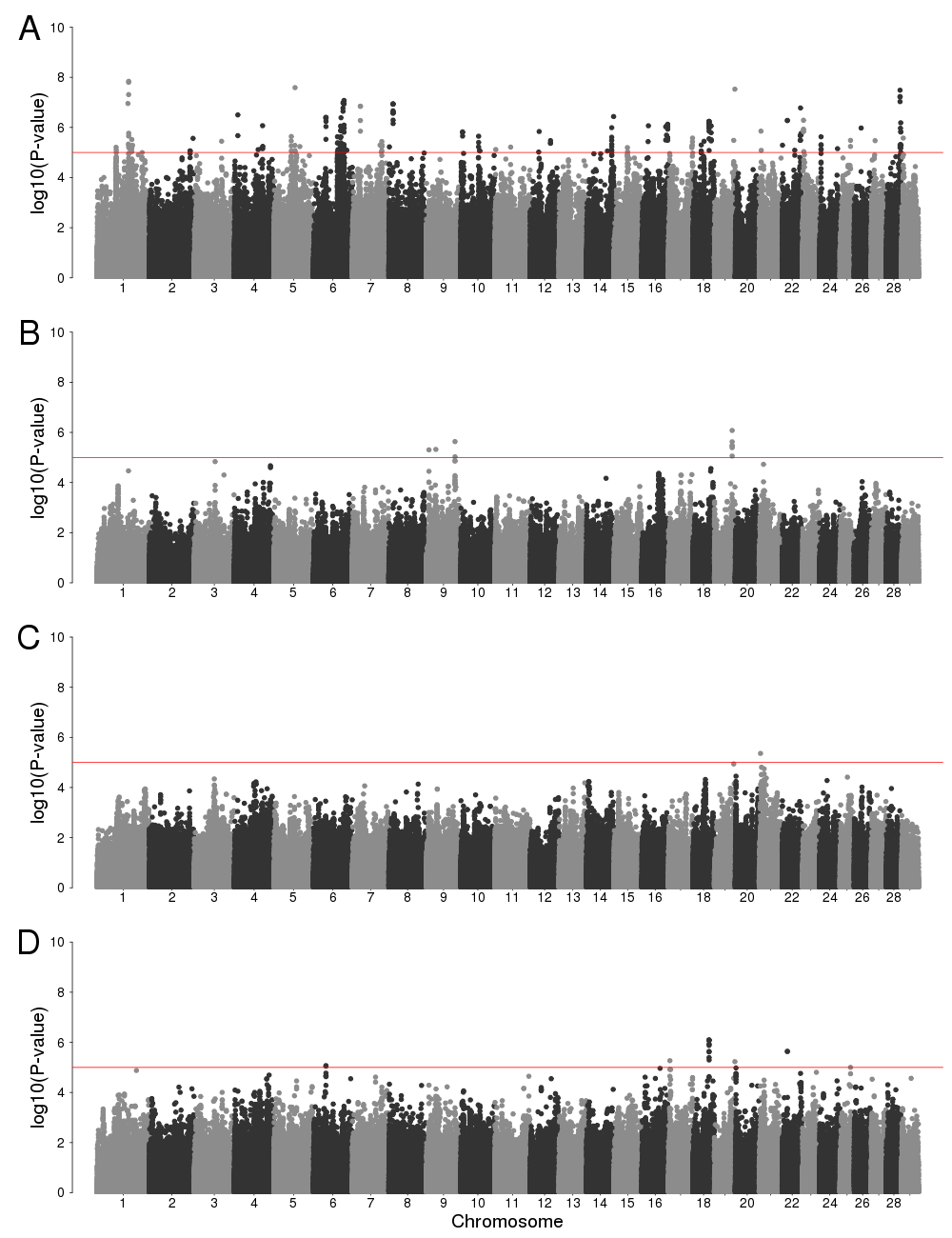
Figure S8 Manhattan plots of genotype-by-environment genome-wide association analysis using maximum vapor pressure deficit as environmental variable for birth weight (*A*), weaning weight (*B*), yearling weight (*C*), and using multivariate analysis (*D*). Horizontal red line indicates a significant threshold (*P* < 1e-5).

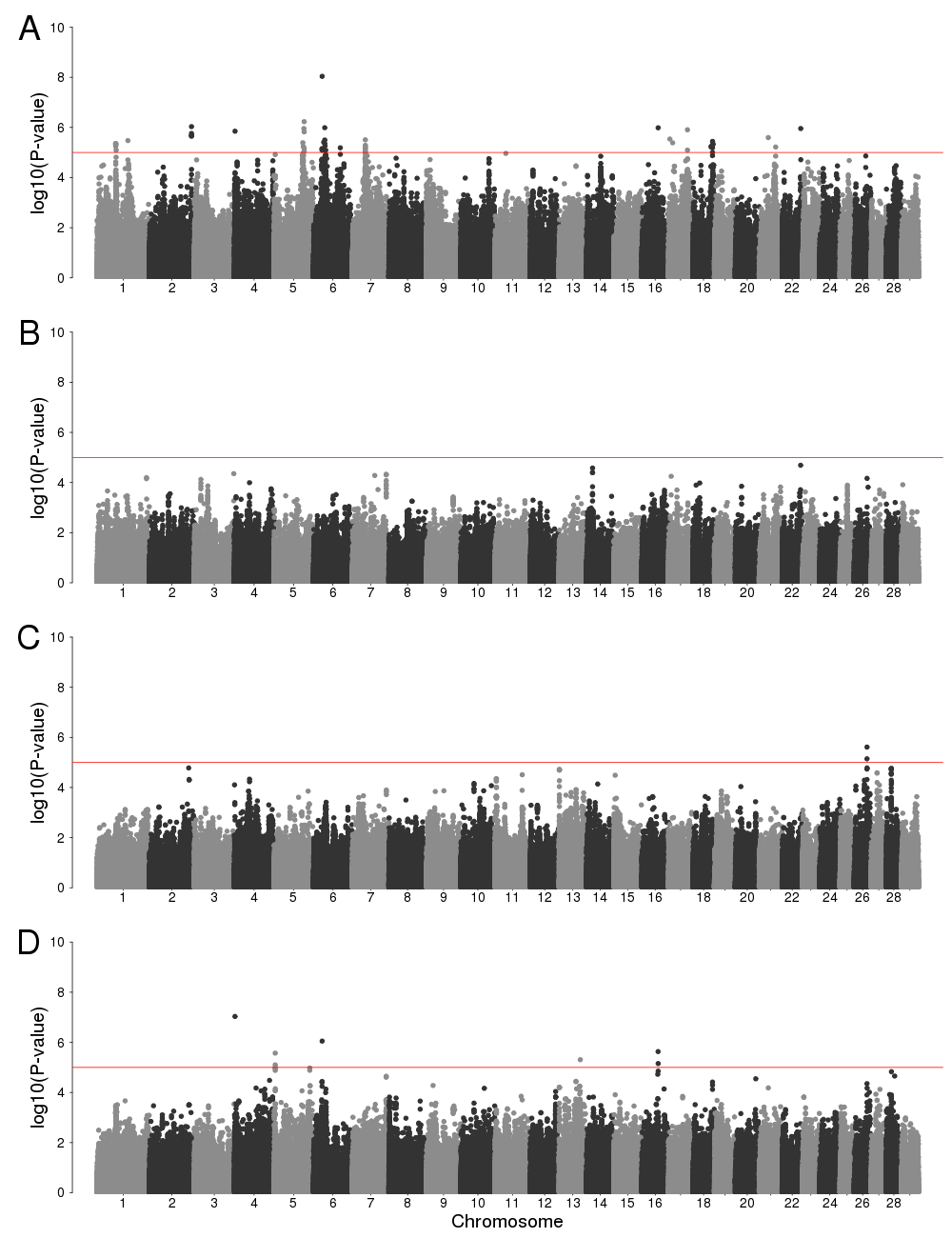
Figure S9 Manhattan plots of genotype-by-environment genome-wide association analysis using Southeast ecoregion as environmental factor for birth weight (*A*), weaning weight (*B*), yearling weight (*C*), and using multivariate analysis (*D*). Horizontal red line indicates a significant threshold (*P* < 1e-5).

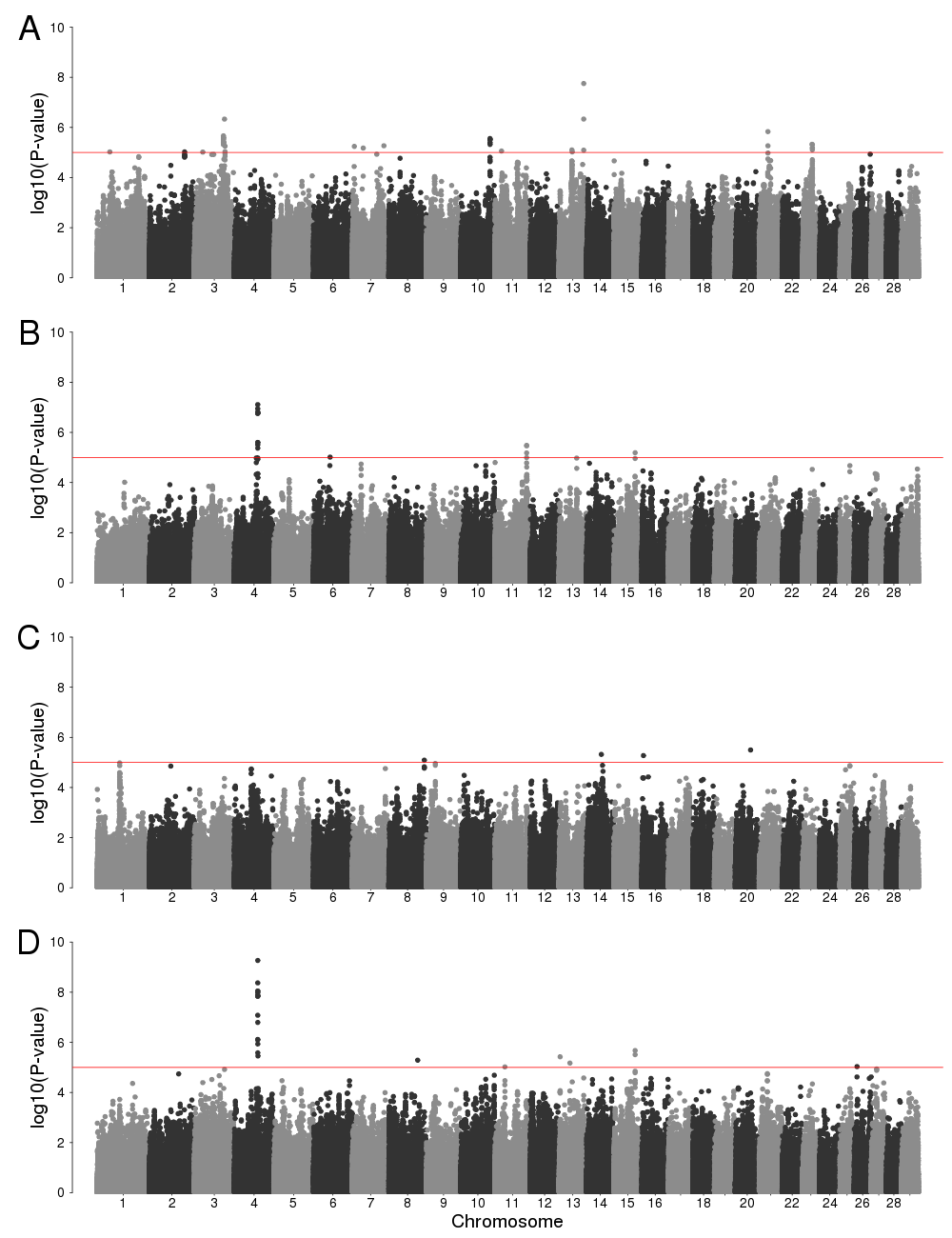
Figure S10 Manhattan plots of genotype-by-environment genome-wide association analysis using High Plains ecoregion as environmental factor for birth weight (*A*), weaning weight (*B*), yearling weight (*C*), and using multivariate analysis (*D*). Horizontal red line indicates a significant threshold (*P* < 1e-5).

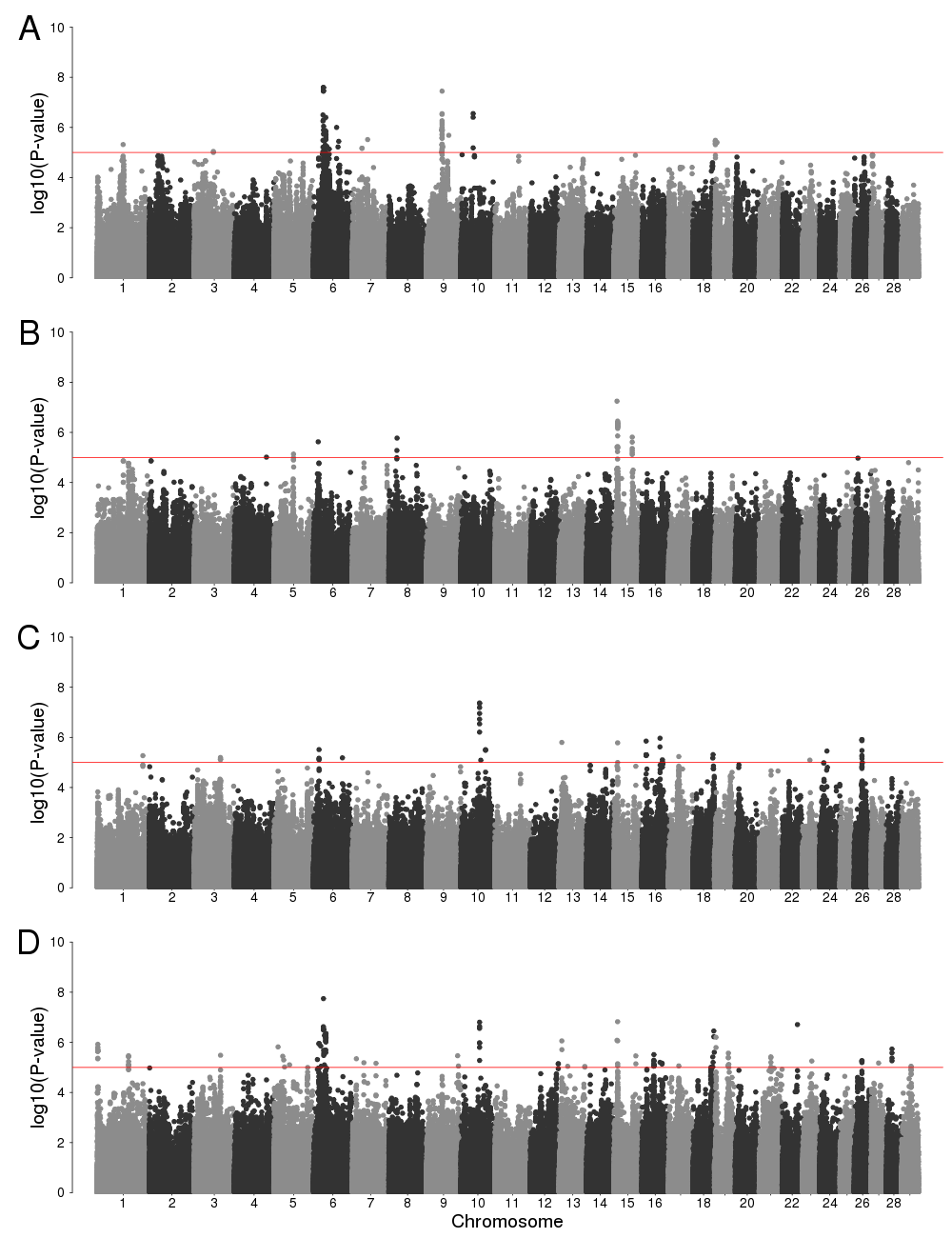
Figure S11 Manhattan plots of genotype-by-environment genome-wide association analysis using Forested Mountains ecoregion as environmental factor for birth weight (*A*), weaning weight (*B*), yearling weight (*C*), and using multivariate analysis (*D*). Horizontal red line indicates a significant threshold (*P* < 1e-5).

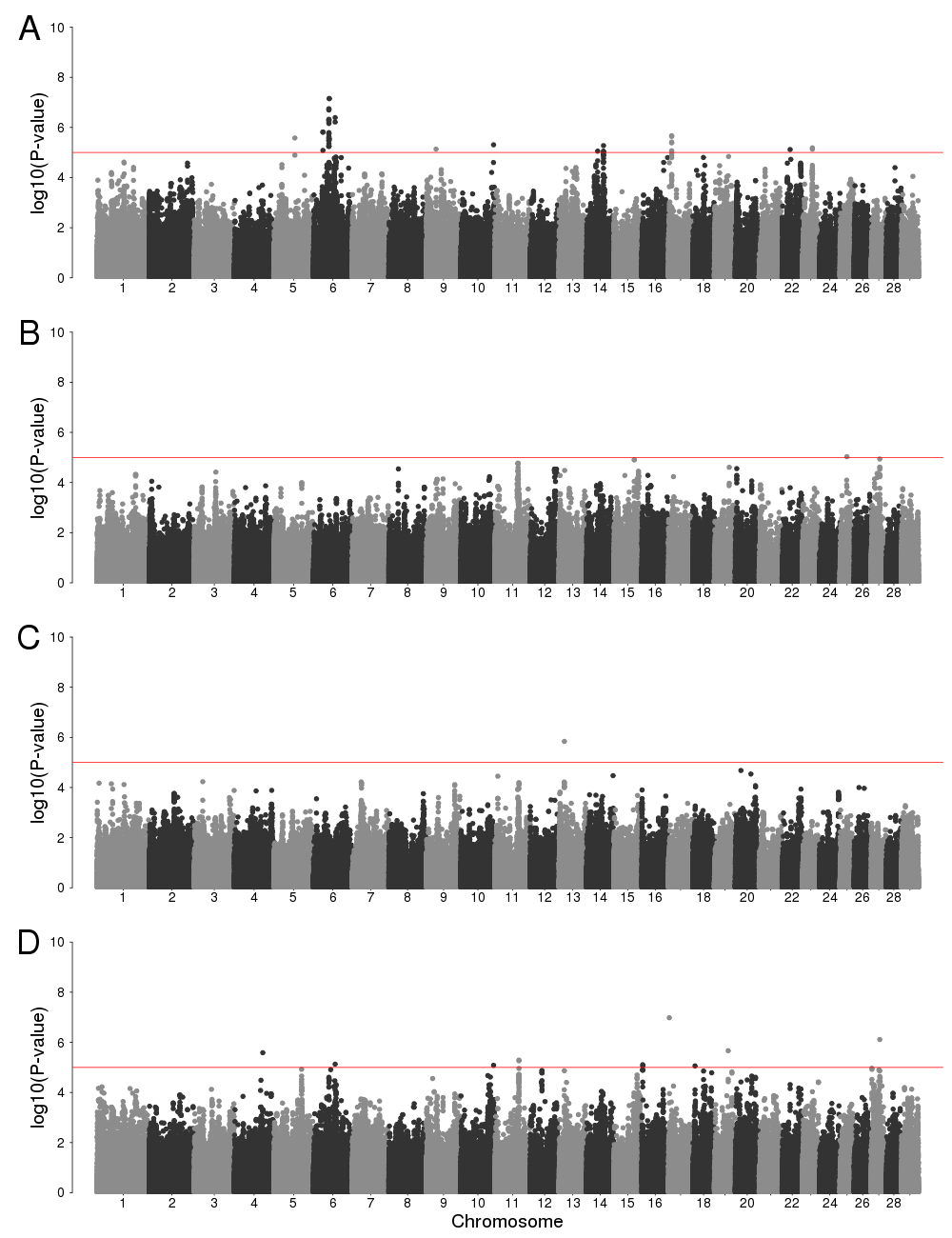
Figure S12 Manhattan plots of genotype-by-environment genome-wide association analysis using Fescue Belt ecoregion as environmental factor for birth weight (*A*), weaning weight (*B*), yearling weight (*C*), and using multivariate analysis (*D*). Horizontal red line indicates a significant threshold (*P* < 1e-5).

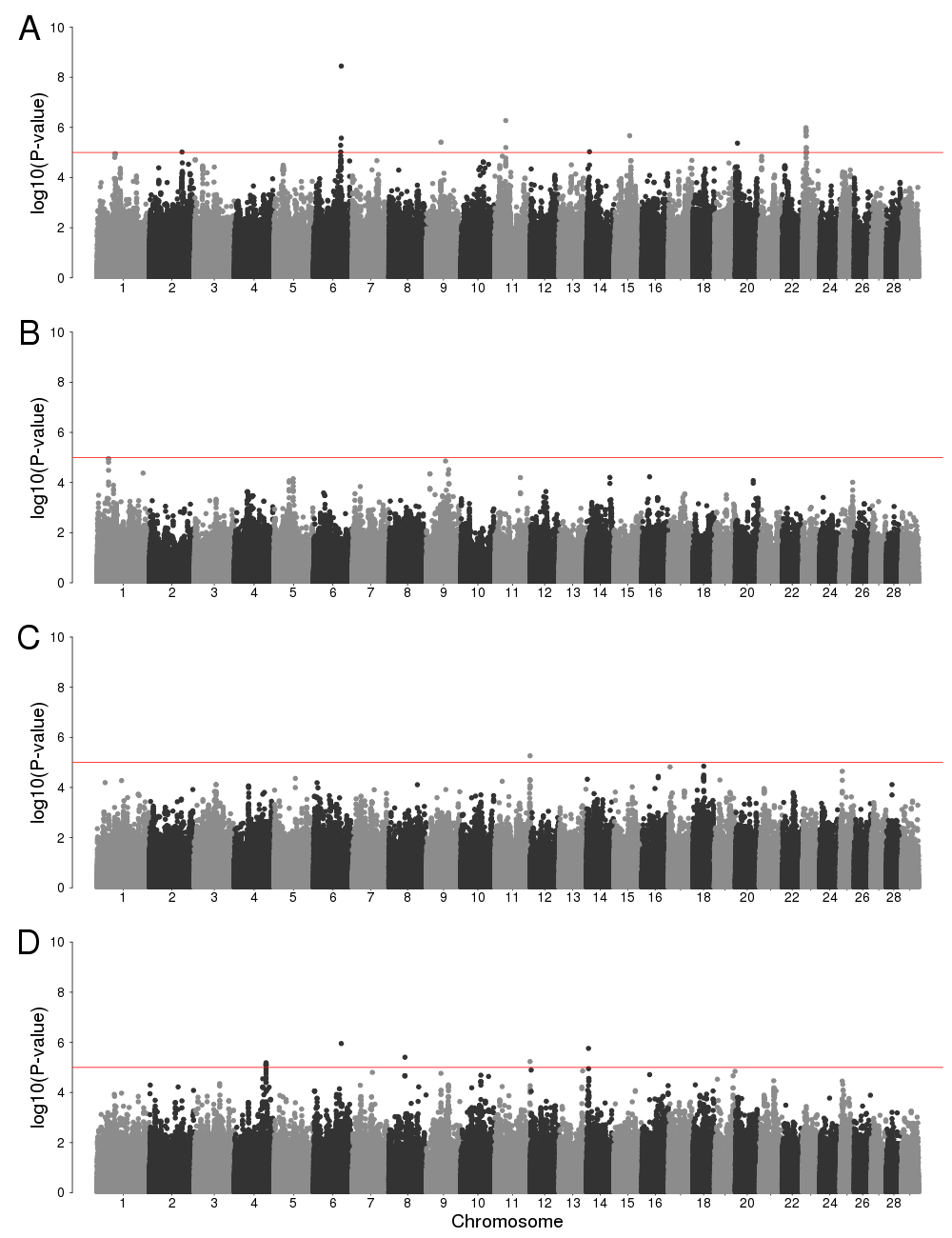
Fig. S13. Manhattan plots of genotype-by-environment genome-wide association analysis using Upper Midwest & Northeast ecoregion as environmental factor for birth weight (*A*), weaning weight (*B*), yearling weight (*C*), and using multivariate analysis (*D*). Horizontal red line indicates a significant threshold (*P* < 1e-5).

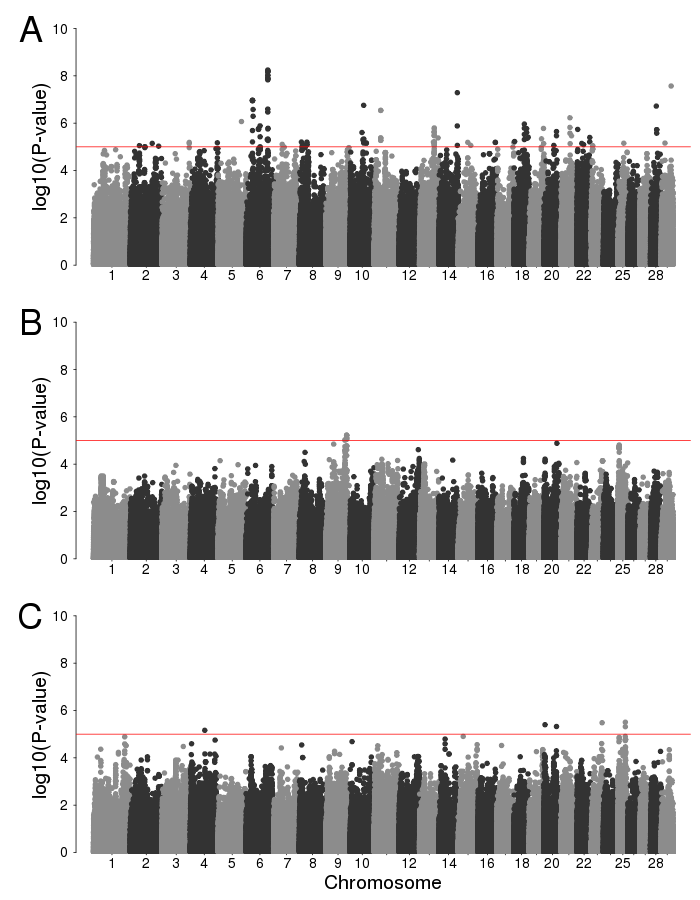
Fig. S14. Manhattan plots of genotype-by-environment genome-wide association analysis using Desert & Arid Prairie ecoregion as environmental factor for birth weight (*A*), weaning weight (*B*), yearling weight (*C*). Horizontal red line indicates a significant threshold (*P* < 1e-5).

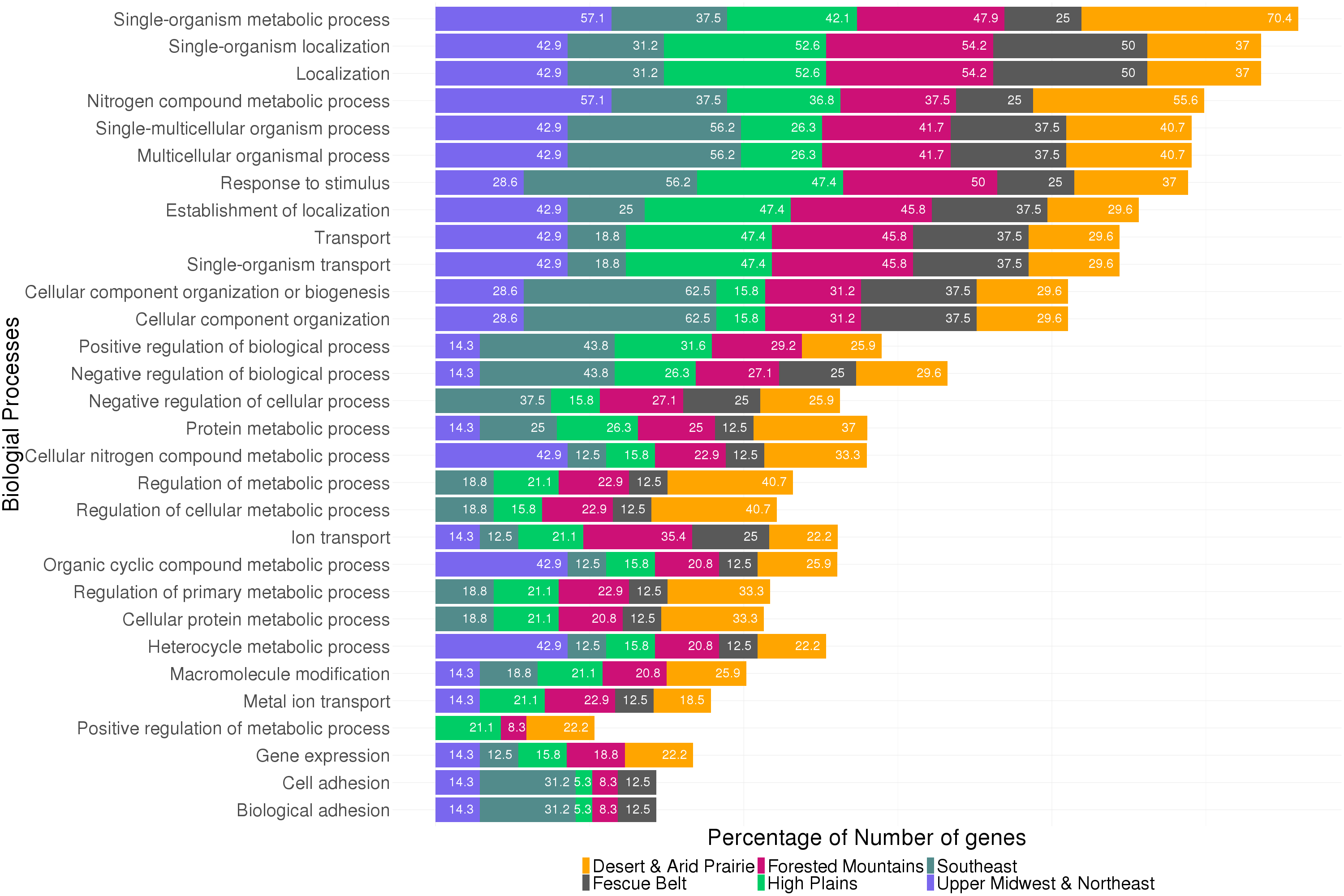
Fig. S15. Percentage of candidate genes with genotype-by-environment effects for each ecoregion that participate in the enriched gene ontology (GO) terms. Genes were located 10 kb from significant SNPs (*P* < 1e-5).

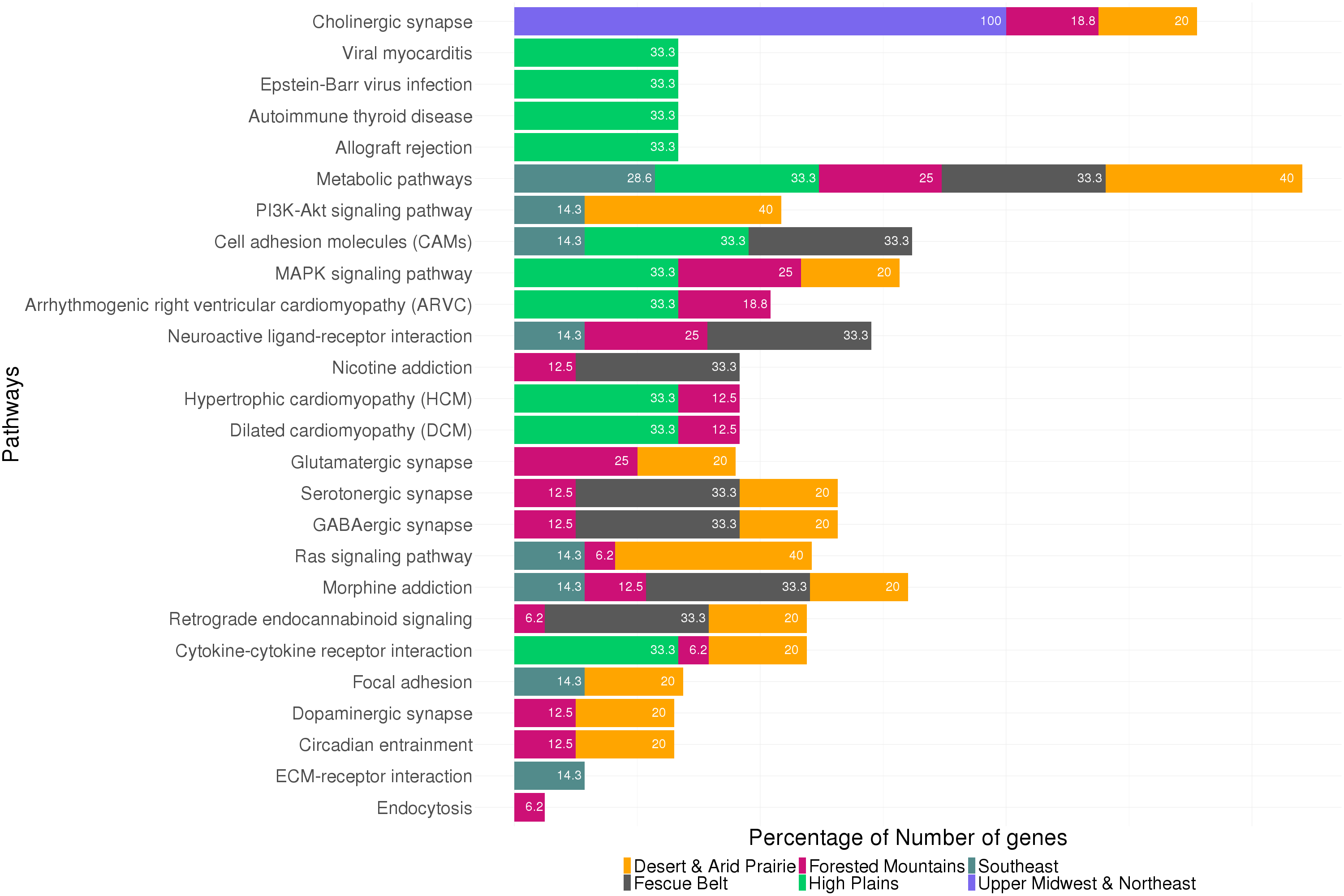
Fig. S16. Percentage of candidate genes with genotype-by-environment effects for each ecoregion that participate in the enriched biological pathways. Genes were located 10 kb from significant SNPs (*P* < 1e-5).

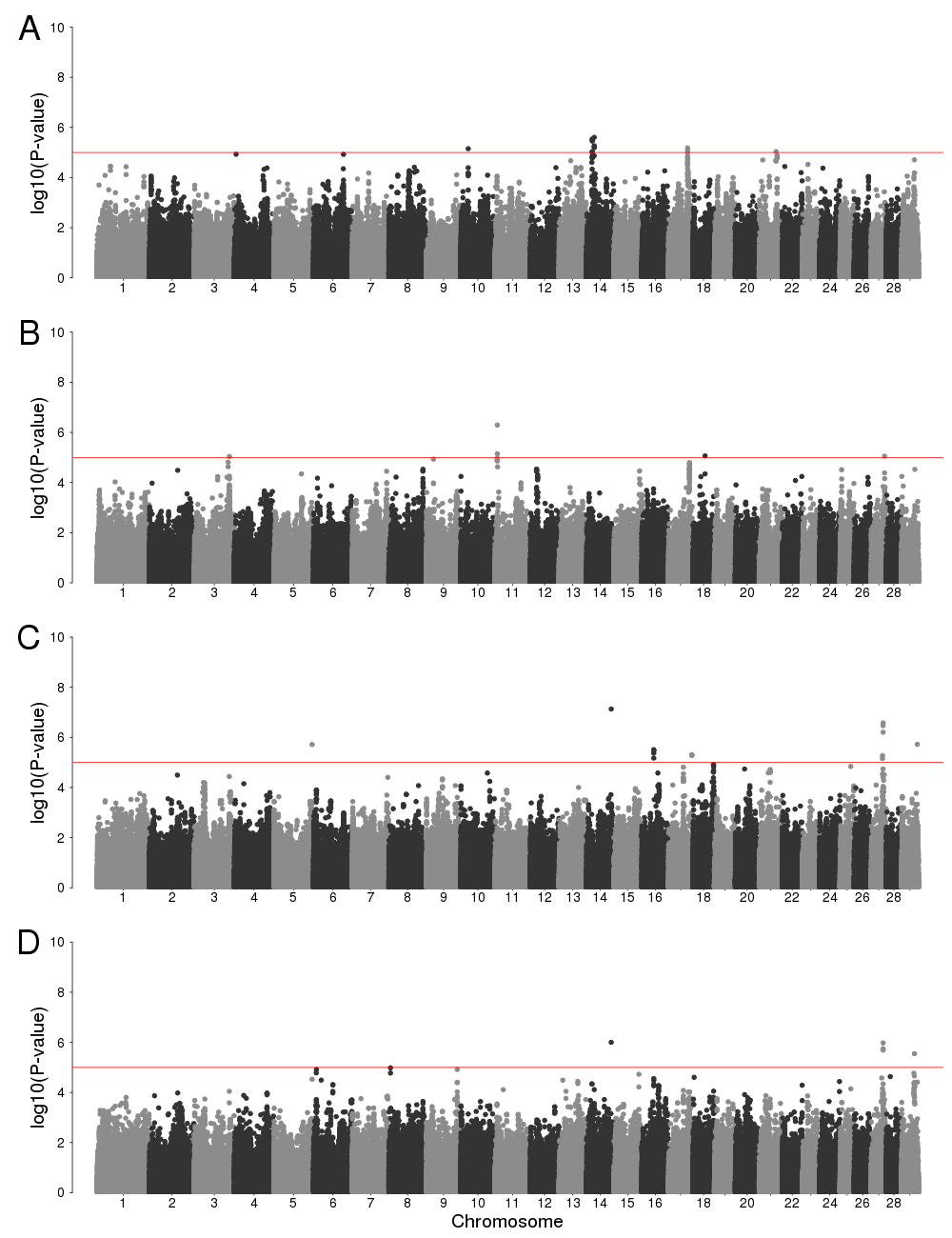
Fig. S17. Manhattan plots of variance-heterogeneity genome-wide association analysis using residuals accounting for only additive effects for birth weight (*A*), weaning weight (*B*), yearling weight (*C*), and using multivariate analysis (*D*). Horizontal red line indicates a significant threshold (*P* < 1e-5).

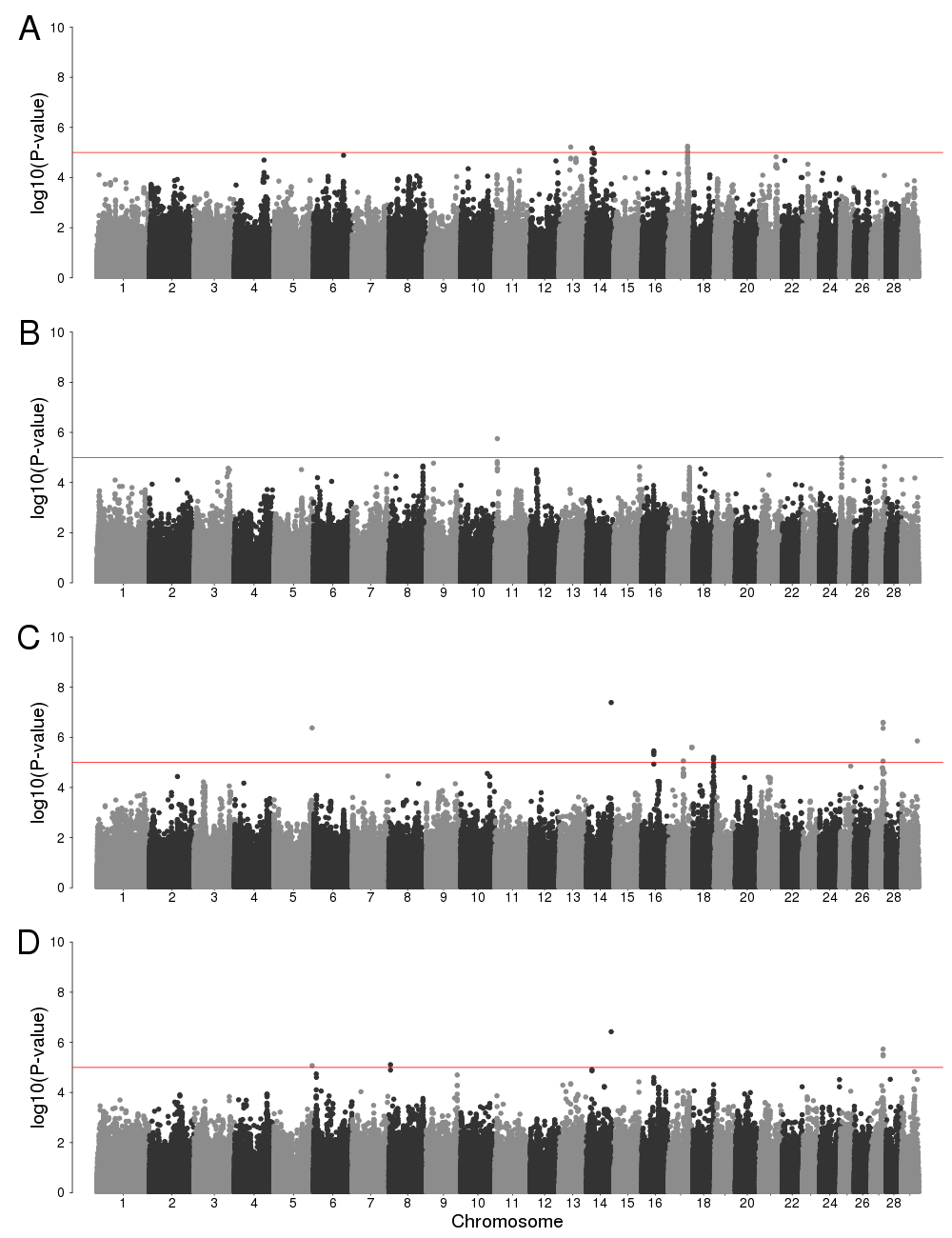
Fig. S18. Manhattan plots of variance-heterogeneity genome-wide association analysis using residuals accounting for additive, dominance and epistatic effects for birth weight (*A*), weaning weight (*B*), yearling weight (*C*), and using multivariate analysis (*D*). Horizontal red line indicates a significant threshold (*P* < 1e-5).

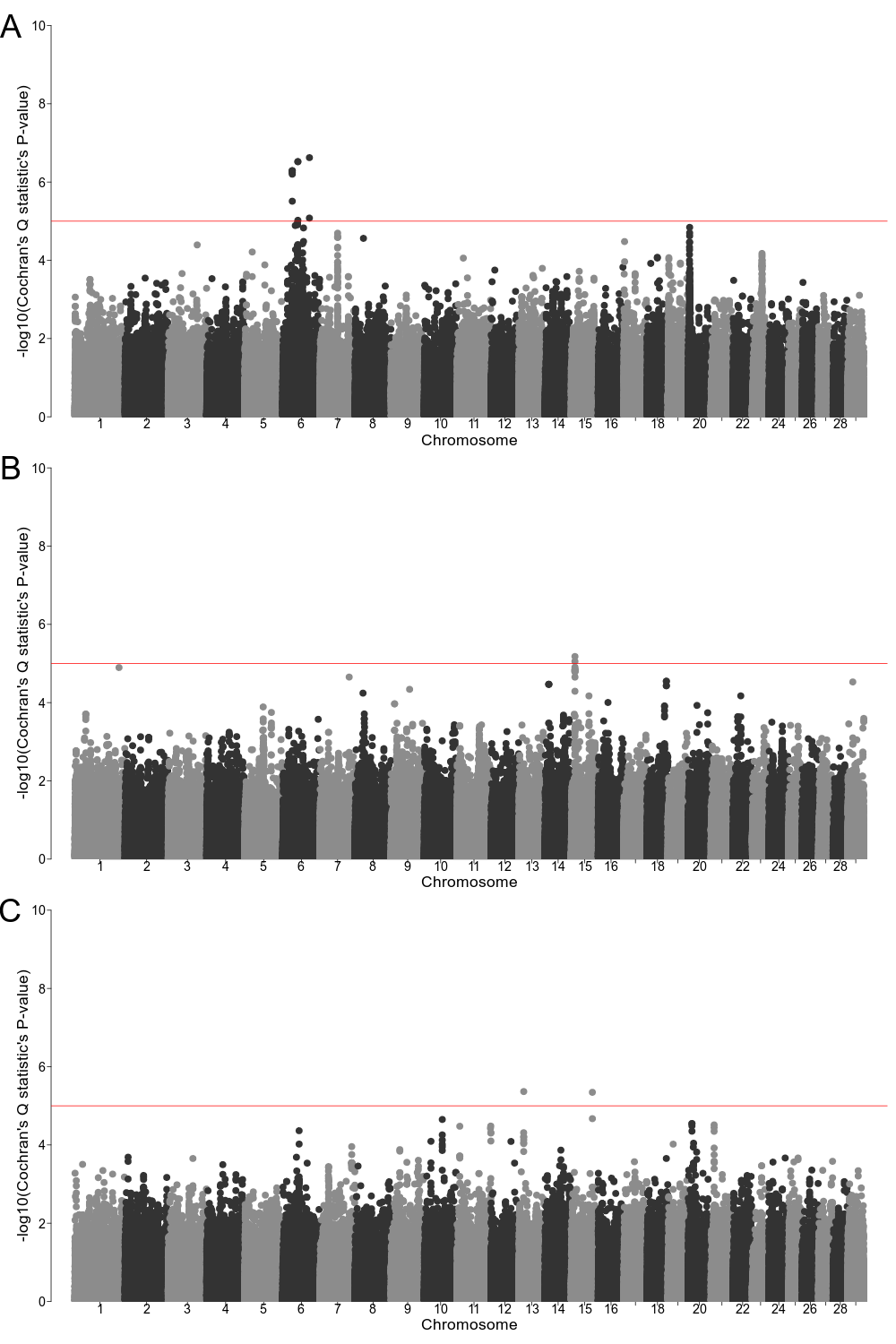
Fig. S19. Meta-analysis of ecoregion-specific genome-wide association analysis. (*A*) Manhattan plot of Cochran's Q statistic's *P*-value for birth weight. (*B*) Manhattan plot of Cochran's Q statistic's *P*-value for weaning weight. (*C*) Manhattan plot of Cochran's Q statistic's *P*-value for yearling weight. Horizontal red line indicates a significant threshold (*P* < 1e-5).

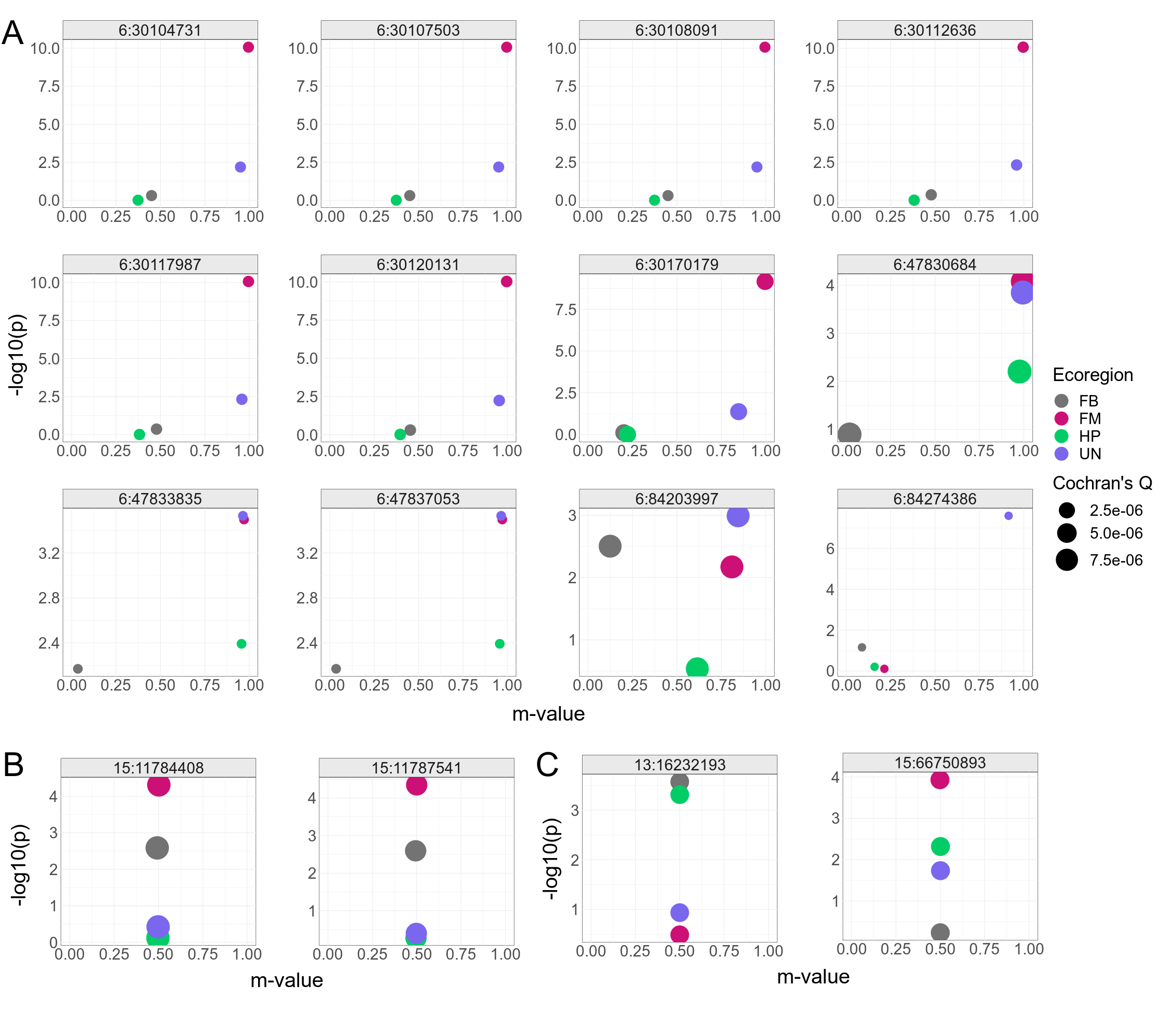
**Fig. S20.** PM-plot (ecoregion-specific *P*-value and the posterior probability of an effect) from meta-analysis of ecoregion-specific GWAA for birth weight (*A*), weaning weight (*B*), and yearling weight (*C*). Points are colored by ecoregion and sized based on Cochran's Q statistic's *P*-value. United States ecoregions were represented as Fescue Belt (FB), Forested Mountains (FM), High Plains (HP), and Upper Midwest & Northeast (UN).

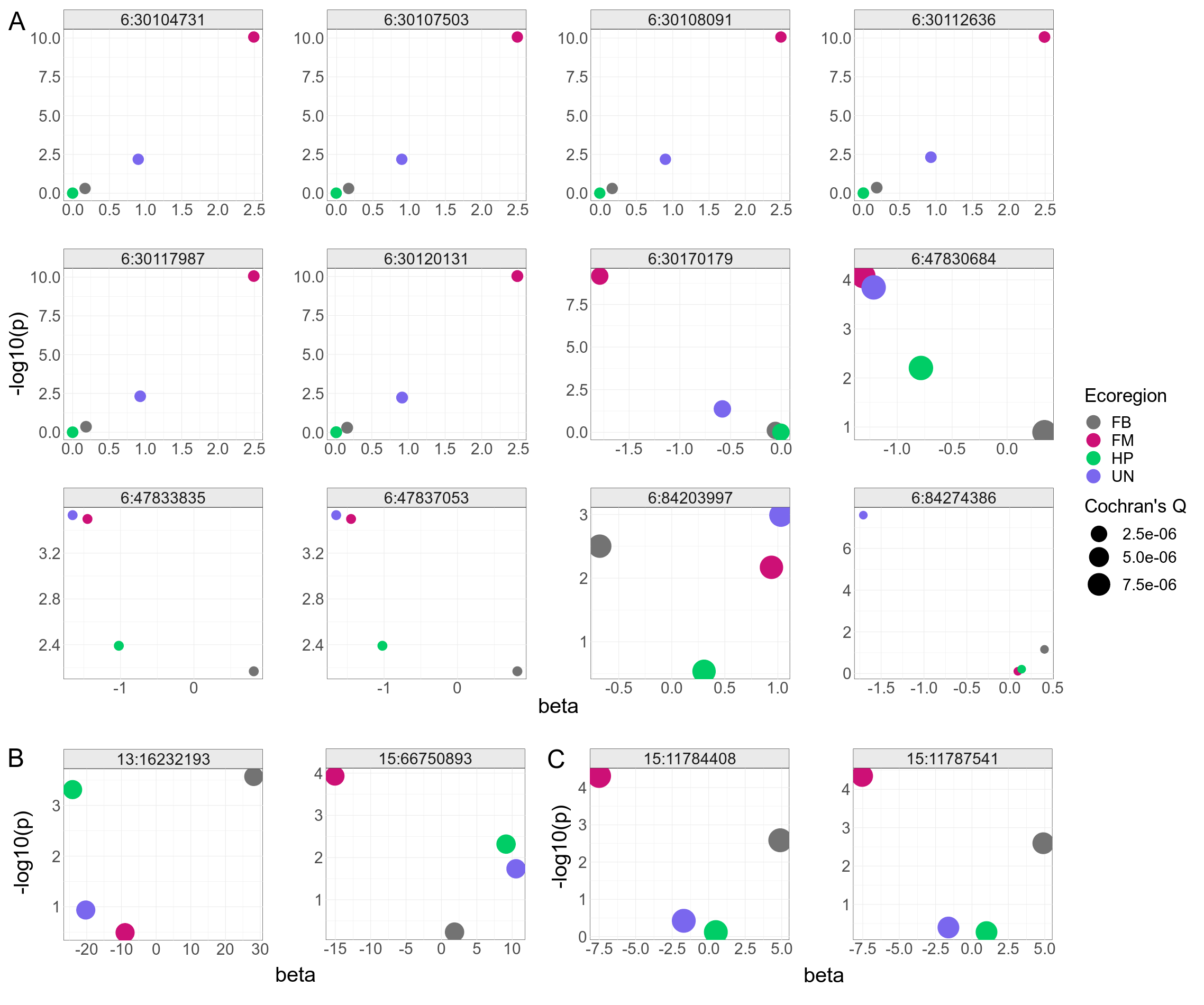
**Fig. 21.** PB-plot (ecoregion-specific *P*-value and the effect size) from meta-analysis of ecoregion-specific GWAA for birth weight (*A*), weaning weight (*B*), and yearling weight (*C*). Points are colored by ecoregion and sized based on Cochran's Q statistic's *P*-value. United States ecoregions were represented as Fescue Belt (FB), Forested Mountains (FM), High Plains (HP), and Upper Midwest & Northeast (UN).

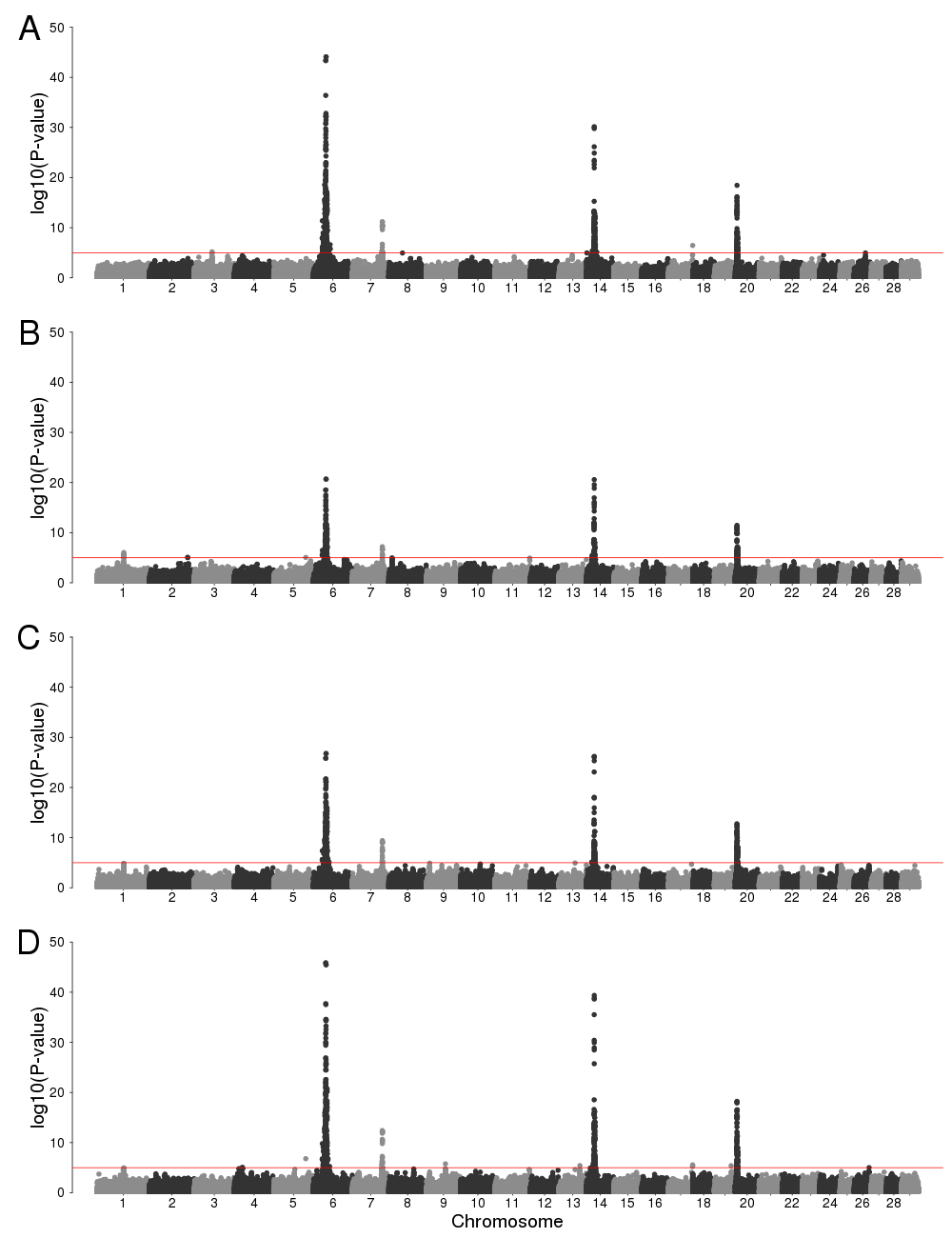
Fig. S22. Manhattan plots of genome-wide association analysis for birth weight (*A*), weaning weight (*B*), yearling weight (*C*), and using multivariate analysis (*D*). Horizontal red line indicates a significant threshold (*P* < 1e-5).

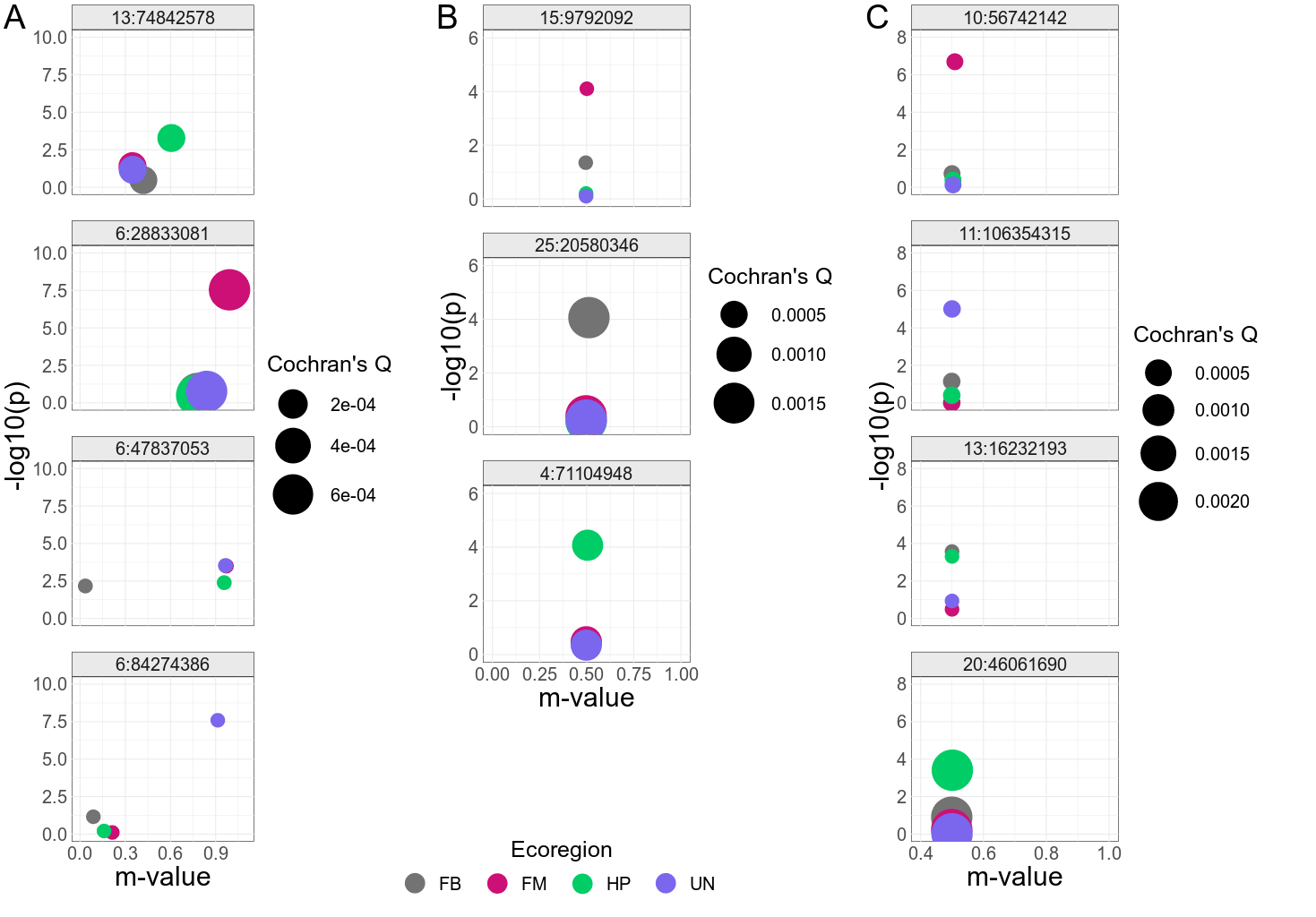
Fig. S23. PM-plot (ecoregion-specific *P*-value and the posterior probability of an effect) from meta-analysis of ecoregion-specific GWAA of some of the most significant GxE SNPs identified by GxE GWAA for birth weight (*A*), weaning weight (*B*), and yearling weight (*C*). Points are colored by ecoregion and sized based on Cochran's Q statistic's *P*-value. United States ecoregions were represented as Fescue Belt (FB), Forested Mountains (FM), High Plains (HP), and Upper Midwest & Northeast (UN). Based on GxE GWAA results, SNPs 15:9792092, 6:28833081, 6:47837053, and 6:84274386 interact with HP, FM, FB, and UN ecoregions, respectively, and affect BW; SNPs 13:74842578, 25:30580346, and 4:71104948 interact with FM, FB, HP ecoregions, respectively, influencing WW; SNPs 10:56742142, 11:106354315, 13:16232193, and 20:46061690 interact with FM, UN, FB, and HP ecoregions, respectively, affecting YW.

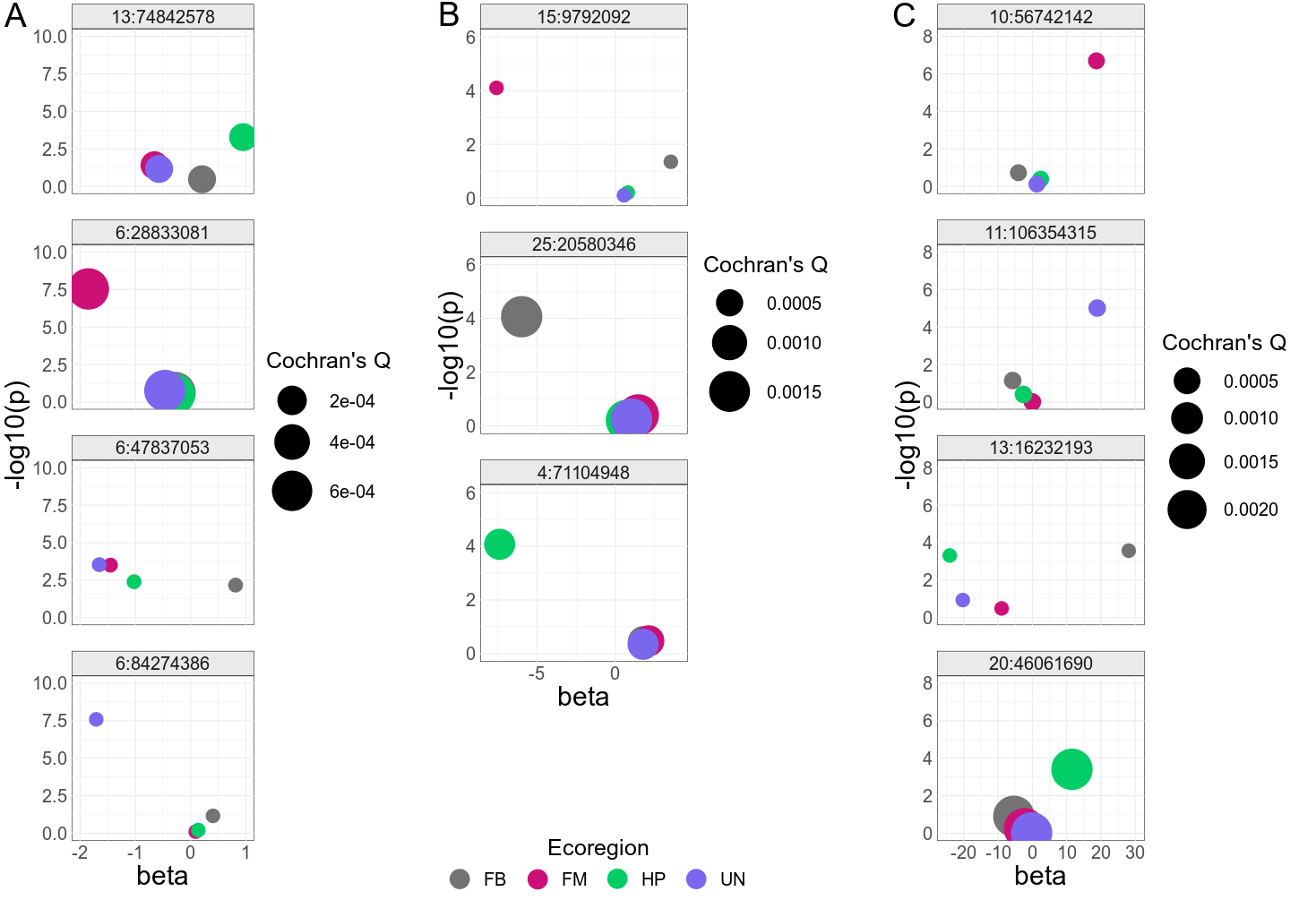
Fig. S24. PB-plot (ecoregion-specific *P*-value and the effect size) from meta-analysis of ecoregion-specific GWAA of some of the most significant GxE SNPs identified by GxE GWAA for birth weight (*A*), weaning weight (*B*), and yearling weight (*C*). Points are colored by ecoregion and sized based on Cochran's Q statistic's *P*-value. United States ecoregions were represented as Fescue Belt (FB), Forested Mountains (FM), High Plains (HP), and Upper Midwest & Northeast (UN). Based on GxE GWAA results, SNPs 15:9792092, 6:28833081, 6:47837053, and 6:84274386 interact with HP, FM, FB, and UN ecoregions, respectively, and affect birth weight; SNPs 13:74842578, 25:30580346, and 4:71104948 interact with FM, FB, HP ecoregions, respectively, influencing weaning weight; SNPs 10:56742142, 11:106354315, 13:16232193, and 20:46061690 interact with FM, UN, FB, and HP ecoregions, respectively, affecting yearling weight.

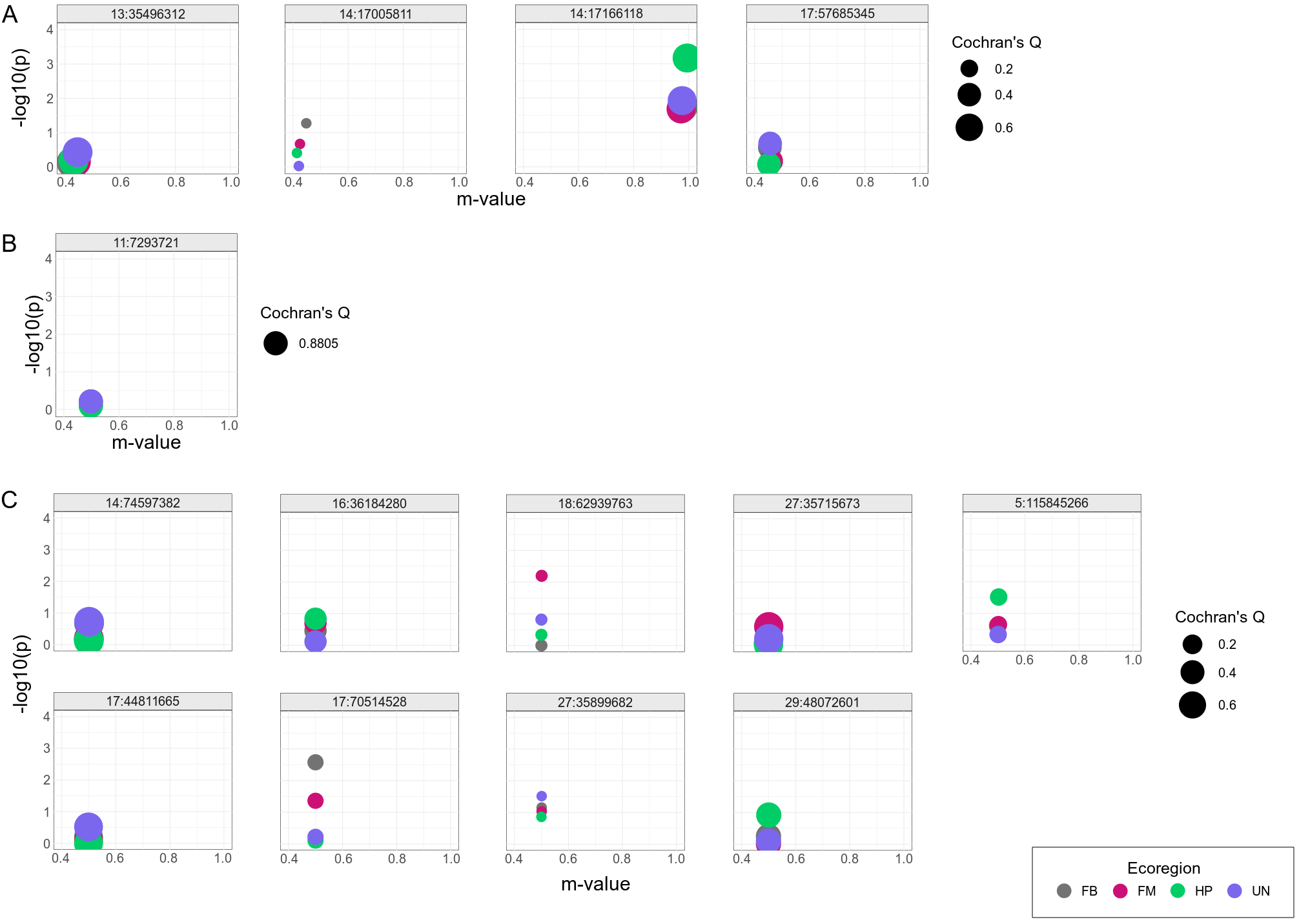
Fig. S25. PM-plot (ecoregion-specific *P*-value and the posterior probability of an effect) from meta-analysis of ecoregion-specific GWAA of variance-heterogeneity SNPs identified by vGWAA for birth weight (*A*), weaning weight (*B*), and yearling weight (*C*). Points are colored by ecoregion and sized based on Cochran's Q statistic's *P*-value. United States ecoregions were represented as Fescue Belt (FB), Forested Mountains (FM), High Plains (HP), and Upper Midwest & Northeast (UN).

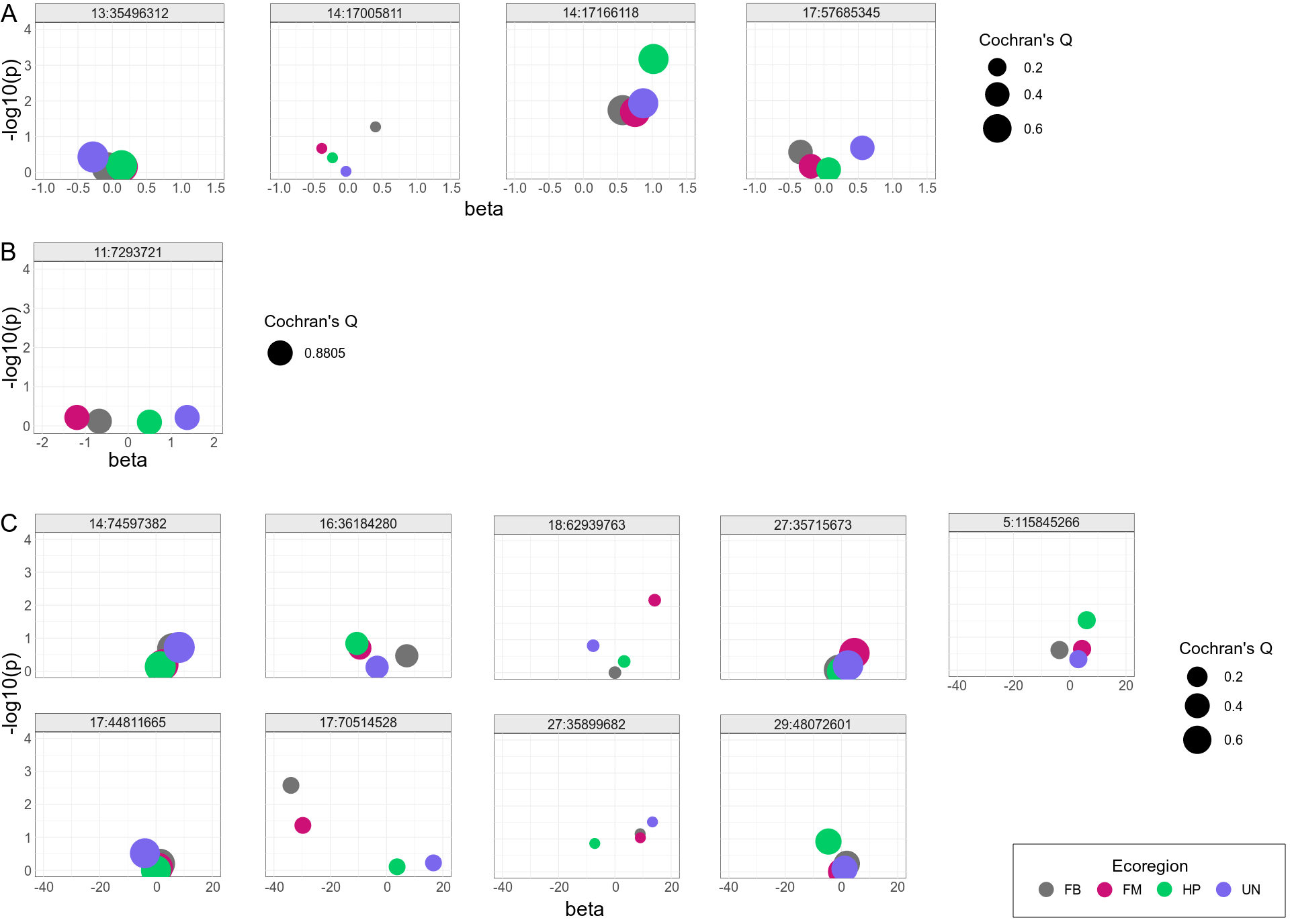
Fig. S26. PB-plot (ecoregion-specific *P*-value and the effect size) from meta-analysis of ecoregion-specific GWAA of variance-heterogeneity SNPs identified by vGWAA for birth weight (*A*), weaning weight (*B*), and yeraling weight (*C*). Points are colored by ecoregion and sized based on Cochran's Q statistic's *P*-value. United States ecoregions were represented as Fescue Belt (FB), Forested Mountains (FM), High Plains (HP), and Upper Midwest & Northeast (UN).
