## Additional File 2 for "Extensive genome-wide association analyses identify genotype-by-environment interactions of growth traits in Simmental cattle"

Supplementary Tables

**Table S1.** Summary statistics for growth traits and environmental variables across the US ecoregions.

| Traits/Env.Variables | SE | HP | FM | FB | UN | DA | Total |
| --- | --- | --- | --- | --- | --- | --- | --- |
| Birth Weight (kg) |  |  |  |  |  |  |  |
| N | 1151 | 3271 | 2397 | 3924 | 2173 | 511 | 13427 |
| Mean | 35.5 | 39.9 | 38.7 | 37.1 | 38.9 | 37.2 | 38.2 |
| SD | 4.4 | 4.9 | 4.8 | 4.4 | 4.5 | 5.2 | 4.8 |
| Weaning Weight (kg) |  |  |  |  |  |  |  |
| N | 873 | 3283 | 2329 | 3025 | 1883 | 454 | 11847 |
| Mean | 303.6 | 313.0 | 287.1 | 296.8 | 315.7 | 289.5 | 302.6 |
| SD | 42.2 | 44.0 | 38.6 | 46.2 | 43.3 | 42.0 | 44.6 |
| Yearling Weight (kg) |  |  |  |  |  |  |  |
| N | 533 | 2525 | 1999 | 1926 | 1293 | 270 | 8546 |
| Mean | 491.5 | 551.4 | 512.0 | 484.9 | 542.5 | 486.0 | 520.1 |
| SD | 78.2 | 81.1 | 71.5 | 89.7 | 81.5 | 71.6 | 85.1 |
| Multivariate |  |  |  |  |  |  |  |
| N | 533 | 2307 | 1992 | 1917 | 1278 | 267 | 8294 |
| Elevation (m) |  |  |  |  |  |  |  |
| Mean | 170.2 | 681.8 | 1587.0 | 341.6 | 372.0 | 914.8 | 747.5 |
| SD | 89.9 | 215.1 | 307.5 | 147.5 | 65.9 | 496.7 | 517.2 |
| Precipitation (ml) |  |  |  |  |  |  |  |
| Mean | 1289.8 | 420.7 | 390.6 | 946 | 749.2 | 497.6 | 643.8 |
| SD | 188.1 | 118.4 | 90.3 | 155.3 | 111.7 | 150.3 | 329.5 |
| Mean Temp. (^o^C) |  |  |  |  |  |  |  |
| Mean | 17.1 | 7.2 | 6.8 | 12.3 | 7.3 | 14.1 | 9.9 |
| SD | 1.6 | 1.8 | 1.4 | 1.3 | 1.4 | 3.9 | 3.8 |
| Min. Temp. (^o^C) |  |  |  |  |  |  |  |
| Mean | 11.1 | 0.6 | -0.8 | 6.1 | 1.8 | 7.4 | 3.3 |
| SD | 1.8 | 1.6 | 1.4 | 1.2 | 1.4 | 4.9 | 4.0 |
| Max. Temp. (^o^C) |  |  |  |  |  |  |  |
| Mean | 23.7 | 14.2 | 14.5 | 18.3 | 13.3 | 22.8 | 16.4 |
| SD | 1.6 | 2.1 | 1.8 | 1.5 | 1.5 | 3.1 | 3.7 |
| Mean Dew point Temp. (^o^C) |  |  |  |  |  |  |  |
| Mean | 11.5 | 0.2 | -2.2 | 6.3 | 2.5 | 5.8 | 3.0 |
| SD | 1.7 | 1.5 | 1.0 | 1.2 | 1.9 | 5.2 | 4.3 |
| Min. Vapor Pres. Deficit (hPs) |  |  |  |  |  |  |  |
| Mean | 1.0 | 1.3 | 1.4 | 1.1 | 0.9 | 1.7 | 1.2 |
| SD | 0.5 | 0.4 | 0.3 | 0.2 | 0.2 | 0.6 | 0.4 |
| Max. Vapor Pres. Deficit (hPs) |  |  |  |  |  |  |  |
| Mean | 17.0 | 13.0 | 13.7 | 13.2 | 10.0 | 21.0 | 13.3 |
| SD | 1.7 | 2.1 | 1.8 | 1.6 | 1.1 | 2.6 | 2.9 |

Number of animals (N), mean (mean) and standard deviation (SD) of the traits used in the univariate (Birth Weight, Weaning Weight, and Yearling Weight) and in the multivariate analysis, along with the environmental variables (Env.Variables) such as elevation, precipitation, mean temperature, minimum (Min) temperature, maximum (Max) temperature, mean dew point temperature, minimum (Min) vapor pressure deficit and maximum (Max) vapor pressure deficit. Those information were provided using the full dataset (Total) and by ecoregion, which were named as Southeast (SE), High Plains (HP), Forested Mountains (FM), Fescue Belt (FB), Upper Midwest & Northeast (UN), Desert & Arid Plains (DA).

**Table S2.** Top 30 enriched gene ontology (GO) terms for candidate genes located at 10 kb and 100 kb sequence window from the significant SNPs (*P* < 1e-05).

| GO Term | GO name | 10 kb window | | 100 kb window | |
| --- | --- | --- | --- | --- | --- |
|  |  | N | *P*-value_adj_ | N | *P*-value_adj_ |
| GO:0050896 | Response to stimulus | 164 | 1.9e-11 | 452 | 5.7e-33 |
| GO:0006807 | Nitrogen compound metabolic process | 178 | 1.4e-08 | 541 | 6.7e-28 |
| GO:0019222 | Regulation of metabolic process | 107 | 1.5e-09 | 335 | 2.2e-27 |
| GO:1901360 | Organic cyclic compound metabolic process | 85 | 4.7e-07 | 294 | 3.4e-27 |
| GO:0032501 | Multicellular organismal process | 150 | 2.3e-07 | 404 | 5.2e-27 |
| GO:0034641 | Cellular nitrogen compound metabolic process | 92 | 3.2e-08 | 311 | 6.7e-27 |
| GO:0044707 | Single-multicellular organism process | 150 | 5.0e-07 | 404 | 1.3e-26 |
| GO:0051716 | Cellular response to stimulus | - | - | 326 | 1.7e-25 |
| GO:0080090 | Regulation of primary metabolic process | 97 | 1.4e-07 | 312 | 1.8e-25 |
| GO:0048518 | Positive regulation of biological process | 105 | 7.6e-10 | 309 | 2.9e-25 |
| GO:0060255 | Regulation of macromolecule metabolic process | - | - | 312 | 1.0e-24 |
| GO:0031323 | Regulation of cellular metabolic process | 98 | 2.3e-07 | 315 | 1.4e-24 |
| GO:0046483 | Heterocycle metabolic process | 81 | 5.7e-07 | 277 | 2.1e-24 |
| GO:0044710 | Single-organism metabolic process | 210 | 1.5e-09 | 618 | 3.7e-24 |
| GO:0006725 | Cellular aromatic compound metabolic process | - | - | 280 | 1.4e-23 |
| GO:0071840 | Cellular component organization or biogenesis | 119 | 9.1e-10 | 280 | 2.4e-22 |
| GO:0051179 | Localization | 128 | 6.6e-10 | 337 | 4.4e-22 |
| GO:0019538 | Protein metabolic process | 118 | 3.7e-09 | 334 | 8.6e-22 |
| GO:0010467 | Gene expression | 77 | 5.4e-07 | 252 | 8.9e-22 |
| GO:0006139 | Nucleobase-containing compound metabolic process | - | - | 271 | 9.7e-22 |
| GO:0032502 | Developmental process | - | - | 339 | 1.9e-21 |
| GO:0044767 | Single-organism developmental process | - | - | 339 | 6.1e-21 |
| GO:0007154 | Cell communication | - | - | 328 | 1.4e-20 |
| GO:0006950 | Response to stress | - | - | 176 | 1.7e-20 |
| GO:0016043 | Cellular component organization | 118 | 1.8e-08 | 338 | 2.1e-20 |
| GO:0007275 | Multicellular organismal development | - | - | 281 | 5.3e-20 |
| GO:0051234 | Establishment of localization | 101 | 6.6e-10 | 264 | 9.5e-20 |
| GO:0048519 | Negative regulation of biological process | 102 | 4.0e-08 | 280 | 2.1e-19 |
| GO:0048856 | Anatomical structure development | - | - | 309 | 3.7e-19 |
| GO:0023052 | Signaling | - | - | 323 | 7.8e-19 |
| GO:1902578 | Single-organism localization | 128 | 1.2e-09 | - | - |
| GO:0006811 | Ion transport | 53 | 2.0e-09 | - | - |
| GO:0006810 | Transport | 97 | 3.8e-09 | - | - |
| GO:0044765 | Single-organism transport | 97 | 3.2e-08 | - | - |
| GO:0048523 | Negative regulation of cellular process | 91 | 3.2e-08 | - | - |
| GO:0007155 | Cell adhesion | 29 | 1.8e-07 | - | - |
| GO:0022610 | Biological adhesion | 30 | 1.8e-07 | - | - |
| GO:0044267 | Cellular protein metabolic process | 95 | 2.1e-07 | - | - |
| GO:0009893 | Positive regulation of metabolic process | 53 | 2.3e-07 | - | - |
| GO:0043412 | Macromolecule modification | 82 | 2.6e-07 | - | - |
| GO:0030001 | Metal ion transport | 30 | 6.8e-07 | - | - |

Number of candidate genes (N); *P*-value adjusted for FDR of 5% (*P*-value_adj_).

**Table S3.** Top 30 enriched pathways for candidate genes located at 10 kb and 100 kb sequence window from the significant SNPs (*P* < 1e-05).

| Pathway entry | Pathway name | 10 kb window | | 100 kb window | |
| --- | --- | --- | --- | --- | --- |
|  |  | N | *P*-value_adj_ | N | *P*-value_adj_ |
| bta01100 | Metabolic pathways | 36 | 1.8e-10 | 82 | 1.5e-15 |
| bta05032 | Morphine addiction | 10 | 2.4e-08 | 18 | 7.3e-12 |
| bta04151 | PI3K-Akt signaling pathway | 11 | 1.4e-03 | 32 | 6.9e-11 |
| bta04723 | Retrograde endocannabinoid signaling | 10 | 4.6e-06 | 18 | 2.2e-09 |
| bta04510 | Focal adhesion | 10 | 1.3e-04 | 21 | 2.5e-09 |
| bta05200 | Pathways in cancer | - | - | 44 | 5.9e-09 |
| bta04727 | GABAergic synapse | 9 | 1.5e-06 | 15 | 8.5e-09 |
| bta05033 | Nicotine addiction | 7 | 1.5e-06 | 11 | 1.2e-08 |
| bta05412 | Arrhythmogenic right ventricular cardiomyopathy (ARVC) | 8 | 3.3e-06 | 13 | 4.2e-08 |
| bta04141 | Protein processing in endoplasmic reticulum | - | - | 18 | 5.8e-08 |
| bta04014 | Ras signaling pathway | 9 | 5.0e-03 | 23 | 8.0e-08 |
| bta03013 | RNA transport | - | - | 20 | 8.0e-08 |
| bta04010 | MAPK signaling pathway | 13 | 5.4e-03 | 29 | 2.9e-07 |
| bta04912 | GnRH signaling pathway | - | - | 12 | 2.4e-06 |
| bta04012 | ErbB signaling pathway | - | - | 12 | 2.9e-06 |
| bta05169 | Epstein-Barr virus infection | 5 | 5.8e-03 | 23 | 6.1e-06 |
| bta04724 | Glutamatergic synapse | 7 | 5.7e-04 | 13 | 7.3e-06 |
| bta04725 | Cholinergic synapse | 6 | 3.2e-03 | 13 | 7.7e-06 |
| bta04921 | Oxytocin signaling pathway | - | - | 15 | 8.7e-06 |
| bta05410 | Hypertrophic cardiomyopathy (HCM) | 7 | 1.0e-04 | 11 | 9.2e-06 |
| bta04726 | Serotonergic synapse | 7 | 1.1e-04 | 13 | 1.0e-05 |
| bta04664 | Fc epsilon RI signaling pathway | - | - | 10 | 1.2e-05 |
| bta04080 | Neuroactive ligand-receptor interaction | 11 | 1.3e-03 | 25 | 1.5e-05 |
| bta05152 | Tuberculosis | - | - | 16 | 2.4e-05 |
| bta05160 | Hepatitis C | - | - | 10 | 2.5e-05 |
| bta04390 | Hippo signaling pathway | - | - | 13 | 2.9e-05 |
| bta04730 | Long-term depression | - | - | 10 | 3.5e-05 |
| bta03030 | DNA replication | - | - | 6 | 6.9e-05 |
| bta05161 | Hepatitis B | - | - | 15 | 6.9e-05 |
| bta04722 | Neurotrophin signaling pathway | - | - | 12 | 6.9e-05 |
| bta05414 | Dilated cardiomyopathy | 7 | 1.3e-04 | - | - |
| bta05416 | Viral myocarditis | 3 | 2.2e-03 | - | - |
| bta04514 | Cell adhesion molecules (CAMs) | 5 | 2.5e-03 | - | - |
| bta04144 | Endocytosis | 9 | 2.5e-03 | - | - |
| bta04976 | Bile secretion | 5 | 3.1e-03 | - | - |
| bta00770 | Pantothenate and CoA biosynthesis | 2 | 3.8e-03 | - | - |
| bta05330 | Allograft rejection | 2 | 3.8e-03 | - | - |
| bta04060 | Cytokine-cytokine receptor interaction | 7 | 3.9e-03 | - | - |
| bta00785 | Lipoic acid metabolism | 2 | 3.9e-03 | - | - |
| bta04728 | Dopaminergic synapse | 6 | 4.7e-03 | - | - |
| bta04512 | ECM-receptor interaction | 2 | 5.0e-03 | - | - |
| bta05320 | Autoimmune thyroid disease | 4 | 5.3e-03 | - | - |
| bta00510 | N-Glycan biosynthesis | 4 | 5.8e-03 | - | - |
| bta04713 | Circadian entrainment | 5 | 6.7e-03 | - | - |

Number of candidate genes (N); *P*-value adjusted for FDR of 5% (*P*-value_adj_).

**Table S4.** Proportion of genetic variance based on the additive genomic model (MA), and additive, dominance and epistasis genomic model (MADE) and their standard errors for each trait.

| Trait | MA | MADE | | |
| --- | --- | --- | --- | --- |
|  | ${\sigma_{A}^{2}}/{\sigma_{P}^{2}}$ | ${\sigma_{A}^{2}}/{\sigma_{P}^{2}}$ | ${\sigma_{D}^{2}}/{\sigma_{P}^{2}}$ | ${\sigma_{Ep}^{2}}/{\sigma_{P}^{2}}$ |
| Birth weight | 0.37 ± 0.01 | 0.33 ± 0.01 | 0.00 ± 0.01 | 0.25 ± 0.03 |
| Weaning weight | 0.30 ± 0.01 | 0.28 ± 0.02 | 0.02 ± 0.02 | 0.12 ± 0.04 |
| Yearling weight | 0.40 ± 0.01 | 0.40 ± 0.02 | 0.05 ± 0.02 | 0.11 ± 0.05 |

${\sigma_{A}^{2}}/{\sigma_{P}^{2}}$**,** ${\sigma_{D}^{2}}/{\sigma_{P}^{2}}$**,** ${\sigma_{Ep}^{2}}/{\sigma_{P}^{2}}$ are proportion of additive, dominance, and epistasis variances to total phenotypic variance, respectively.

**Table S5.** Significant vQTL (*P* < 1e-5) for growth traits when using residuals adjusted for additive effects (MA) or for additive, dominance and epistasis effects (MADE).

| Trait | SNP | BTA | Position | Freq | MA | | MADE | | Genes |
| --- | --- | --- | --- | --- | --- | --- | --- | --- | --- |
|  |  |  |  |  | Effect (kg) | *P*-value | Effect (kg) | *P*-value |  |
| BW | *rs110323698* | 10 | 22,312,496 | 0.127 | 0.026 ± 1.6e-04 | 7.07e-06 | - | - | *DAD1, ABHD4* |
|  | *rs137529355* | 13 | 35,496,312 | 0.278 | - | - | -0.020 ± 9.3e-05 | 6.04e-06 | *LYZL1* |
|  | *rs133623280* | 14 | 17,005,811 | 0.458 | -0.019 ± 7.9e-05 | 2.94e-06 | -0.018 ± 7.9e-05 | 6.69e-06 | *ZHX2* |
|  | *rs109136768* | 14 | 17,007,761 | 0.418 | 0.018 ± 8.6e-05 | 9.60e-06 | - | - |  |
|  | *rs43026996* | 14 | 17,166,118 | 0.344 | 0.022 ± 1.1e-04 | 3.51e-06 | 0.021 ± 1.1e-04 | 6.67e-06 |  |
|  | *rs43186199* | 14 | 17,183,088 | 0.347 | 0.022 ± 1.1e-04 | 3.10e-06 | 0.021 ± 1.1e-04 |  |  |
|  | *rs42796317* | 14 | 17,195,903 | 0.356 | 0.021 ± 1.1e-04 | 9.31e-06 | - | - |  |
|  | *rs109515648* | 14 | 23,128,784 | 0.376 | 0.024 ± 1.4e-04 | 6.04e-06 | - | - | *TMEM68, TGS1, LYN, RPS20, MOS, PLAG1, CHCHD7, SDR16C5* |
|  | *rs137303549* | 14 | 23,298,062 | 0.388 | 0.024 ± 1.4e-04 | 6.48e-06 | - | - |  |
|  | *rs134215421* | 14 | 23,329,375 | 0.385 | 0.025 ± 1.4e-04 | 2.49e-06 | - | - |  |
|  | *rs109815800* | 14 | 23,338,890 | 0.384 | 0.024 ± 1.4e-04 | 5.38e-06 | - | - |  |
|  | *rs41843599* | 17 | 57,685,345 | 0.144 | - | - | 0.026 ± 1.7e-04 | 6.42e-06 | *NOS1, KSR2, FBXO21, TESC, FBXW8* |
|  | *rs41843601* | 17 | 57,685,935 | 0.146 | 0.026 ± 1.6e-04 | 6.60e-06 | 0.026 ± 1.6e-04 | 5.62e-06 |  |
|  | *rs109209365* | 17 | 57,699,496 | 0.141 | - | - | 0.026 ± 1.7e-04 | 8.79e-06 |  |
|  | *rs41847363* | 17 | 57,975,119 | 0.146 | 0.026 ± 1.6e-04 | 7.91e-06 | 0.026 ± 1.6e-04 | 9.43e-06 |  |
|  | *rs134957935* | 21 | 51,465,082 | 0.188 | -0.022 ± 1.2e-04 | 9.21e-06 | - | - | *LRFN5* |
| WW | *rs109967828* | 3 | 106,206,442 | 0.169 | 0.025 ± 1.6e-04 | 9.18e-06 | - | - | *MFSD2A, MYCL, TRIT1, BMP8B* |
|  | *rs43657596* | 11 | 7,293,721 | 0.133 | 0.027 ± 1.8e-04 | 7.17e-06 | - | - | *IL18RAP, SLC9A4, SLC9A2* |
|  | *rs134174267* | 11 | 7,304,992 | 0.101 | 0.035 ± 2.4e-04 | 5.08e-07 | 0.033 ± 2.4e-04 | 1.78e-06 |  |
|  | *rs109287854* | 18 | 37,372,422 | 0.219 | 0.022 ± 1.2e-04 | 8.65e-06 | - | - | *-* |
|  | *rs110803856* | 27 | 40,193,292 | 0.209 | 0.023 ± 1.3e-04 | 8.88e-06 | - | - | *TOP2B, RARB* |
| YW | *rs135719485^£^* | 5 | 115,845,266 | 0.336 | -0.025 ± 1.3e-04 | 1.93e-06 | -0.026 ± 1.3e-04 | 4.19e-07 | *RIBC2, FBLN1, SMC1B* |
|  | *rs133130907^¥^* | 14 | 74,597,382 | 0.092 | 0.044 ± 3.1e-04 | 7.42e-08 | 0.044 ± 3.2e-04 | 4.08e-08 | *MMP16* |
|  | *rs41802941* | 16 | 36,184,280 | 0.037 | 0.057 ± 7.7e-04 | 6.73e-06 | - | - | *XCL2, XCL1, DPT* |
|  | *rs41802957* | 16 | 36,200,777 | 0.050 | 0.050 ± 5.8e-04 | 4.27e-06 | 0.050 ± 5.8e-04 | 4.86e-06 |  |
|  | *rs137574197* | 16 | 36,231,107 | 0.050 | 0.051 ± 5.9e-04 | 3.36e-06 | 0.051 ± 5.9e-04 | 4.02e-06 |  |
|  | *rs109835794* | 16 | 36,242,339 | 0.052 | 0.050 ± 5.6e-04 | 3.09e-06 | 0.050 ± 5.7e-04 | 3.46e-06 |  |
|  | *rs134575841* | 17 | 44,811,665 | 0.432 | - | - | 0.022 ± 1.2e-04 | 8.59e-06 | *GALNT9* |
|  | *rs111011858* | 17 | 70,514,528 | 0.010 | 0.103 ± 2.5e-03 | 5.18e-06 | 0.107 ± 2.5e-03 | 2.53e-06 | *YWHAH, SLC5A1, DEPDC5, SLC5A4* |
|  | *rs110473197* | 17 | 70,523,020 | 0.010 | 0.103 ± 2.5e-03 | 5.08e-06 | 0.107 ± 2.5e-03 | 2.50e-06 |  |
|  | *rs109010785* | 17 | 70,527,754 | 0.010 | 0.103 ± 2.5e-03 | 5.08e-06 | 0.107 ± 2.5e-03 | 2.50e-06 |  |
|  | *rs108982594* | 17 | 70,540,863 | 0.010 | 0.103 ± 2.5e-03 | 5.08e-06 | 0.107 ± 2.5e-03 | 2.50e-06 |  |
|  | *rs378352090* | 18 | 62,939,763 | 0.149 | - | - | 0.034 ± 2.8e-04 | 7.85e-06 | *TTYH1, LENG8, LENG9, LAIR1, CDC42EP5, RPS9, TSEN34, MBOAT7, TMC4, LENG1, CNOT3, PRPF31, TFPT, NDUFA3, OSCAR, TARM1* |
|  | *rs383417308* | 18 | 63,229,878 | 0.155 | - | - | 0.034 ± 2.8e-04 | 7.42e-06 |  |
|  | *rs433152283* | 18 | 63,230,047 | 0.155 | - | - | 0.034 ± 2.8e-04 | 6.22e-06 |  |
|  | *rs43727853* | 27 | 33,796,650 | 0.460 | -0.023 ± 1.3e-04 | 7.00e-06 | - | - | *TACC1* |
|  | *rs110205571* | 27 | 34,027,900 | 0.281 | 0.024 ± 1.4e-04 | 5.33e-06 | - | - | *PLEKHA2* |
|  | *rs110909628* | 27 | 35,715,673 | 0.223 | - | - | 0.026 ± 1.6e-04 | 8.82e-06 | *ZMAT4* |
|  | *rs137312252^*^* | 27 | 35,899,682 | 0.097 | 0.041 ± 3.2e-04 | 3.42e-07 | 0.042 ± 3.2e-04 | 2.68e-07 | *ZMAT4* |
|  | *rs135719206^*^* | 27 | 35,902,981 | 0.092 | 0.042 ± 3.4e-04 | 6.24e-07 | 0.042 ± 3.4e-04 | 4.34e-07 |  |
|  | *rs134473626^*^* | 27 | 35,906,532 | 0.095 | 0.042 ± 3.3e-04 | 2.64e-07 | 0.042 ± 3.3e-04 | 2.51e-07 |  |
|  | *rs136265144* | 29 | 48,072,601 | 0.254 | -0.027 ± 1.5e-04 | 1.90e-06 | -0.027 ± 1.5e-04 | 1.40e-06 | *SHANK2* |
| MV | *rs135719485^£^* | 5 | 115,845,266 | 0.334 | - | - | - | 8.47e-06 | *RIBC2, FBLN1, SMC1B* |
|  | *rs109142386* | 8 | 4,822,842 | 0.258 | - | - | - | 7.78e-06 | *GALNTL6* |
|  | *rs133130907^¥^* | 14 | 74,597,382 | 0.091 | - | 9.98e-07 | - | 8.47e-06 | *MMP16* |
|  | *rs137312252^*^* | 27 | 35,899,682 | 0.098 | - | 2.04e-06 | - | 3.77e-07 | *ZMAT4* |
|  | *rs135719206^*^* | 27 | 35,902,981 | 0.092 | - | 1.06e-06 | - | 3.45e-06 |  |
|  | *rs134473626^*^* | 27 | 35,906,532 | 0.096 | - | 1.81e-06 | - | 3.10e-06 |  |
|  | *rs135379559* | 29 | 39,441,489 | 0.099 | - | 2.83e-06 | - | - | *PAG6, PAG11, LOC528815* |

Birth weight (BW), Weaning weight (WW), Yearling weight (YW), multivariate (MV), allele frequency (Freq), allele substitution effect (Effect).

Table S6. False discovery rates based on *P*-value = 1e-5.

| Analysis | BW | WW | YW | Multivariate |
| --- | --- | --- | --- | --- |
| GWA | 0.0072 | 0.0143 | 0.0097 | 0.0074 |
| vGWA |  |  |  |  |
| MA | 0.5463 | 1.4181 | 0.4417 | 1.4119 |
| MADE | 0.8877 | 7.0906 | 0.3926 | 1.1765 |
| GxE GWA |  |  |  |  |
| Mean Temperature | 0.0128 | 7.0906 | 1.7669 | 0.1023 |
| Minimum Temperature | 0.0094 | 1.1817 | - | 0.0802 |
| Maximum Temperature | 0.0174 | 7.0906 | 1.7669 | 0.3069 |
| Mean Dew Point Temperature | 0.0075 | 1.7726 | 7.0678 | 0.0751 |
| Elevation | 0.0500 | 0.2532 | 0.4711 | 0.1412 |
| Precipitation | 0.0210 | 1.7726 | 7.0678 | 0.5883 |
| Minimum Vapor Pressure Deficit | 0.0467 | 3.5452 | 7.0678 | 0.7059 |
| Maximum Vapor Pressure Deficit | 0.0317 | 0.7090 | 7.0678 | 0.4152 |
| Desert & Arid Prairie | 0.0568 | 0.8863 | 1.1779 | - |
| Southeast | 0.1315 | - | 3.5339 | 1.0085 |
| High Plains | 0.1775 | 0.4431 | 1.7669 | 0.3069 |
| Forested Mountains | 0.0789 | 0.2287 | 0.1442 | 0.0477 |
| Fescue Belt | 0.1511 | 7.0906 | 7.0678 | 0.6418 |
| Upper Midwest & Northeast | 0.2959 | - | 7.0678 | 0.7059 |

Birth weight (BW), Weaning Weight (WW), Yearling Weight (YW), Genome-Wide Association (GWA), variance-heterogeneity GWA (vGWA), Genotype-by-environment interaction GWA (GxE GWA).
